# An intrinsic oscillator underlies visual navigation in ants

**DOI:** 10.1101/2022.04.22.489150

**Authors:** Leo Clement, Sebastian Schwarz, Antoine Wystrach

**Affiliations:** Centre de Recherches sur la Cognition Animale, CNRS, Université Paul Sabatier, Toulouse 31062 cedex 09, France

**Author notes:** Address of correspondence: Léo Clément, Université Paul Sabatier, Centre de Recherches sur la Cognition Animale, CNRS, 31062 Toulouse.

**Keywords:** insect navigation, time series analysis, intrinsic oscillator, lateral accessory lobe

## Abstract

Controlling behavior implies a constant balance between exploration – to gather information – and exploitation – to use this information to reach one’s goal. However, how this tradeoff is achieved in navigating animals is unclear. Here we recorded the paths of two phylogenetically distant visually navigating ant species (*Myrmecia croslandi* and *Iridomyrmex purpureus*) using a trackball-treadmill directly in their habitat. We show that both species continuously produce regular lateral oscillations with bursts of forward movement when facing the general direction of travel, providing a remarkable tradeoff between visual exploration across directions and movement areas. This dynamical signature is conserved across navigational contexts but requires certain visual cues to be fully expressed. Rotational feedback regulates the extent of turns, but is not required to produce them, indicating that oscillations are generated intrinsically. Learnt visual information modulates the oscillation’s amplitudes to fit the task at hand in a continuous manner: an unfamiliar panorama enhances the amplitude of oscillations in both naïve and experienced ants, favoring visual exploration; while a learnt familiar panorama reduces them, favoring exploitation through. The observed dynamical signature readily emerges from a simple neural-circuit model of the insect’s conserved pre-motor area known as the lateral accessory lobe, endorsing oscillations as a core, ancestral way of moving in insects. We discuss the importance and evolution of self-generated behaviors and how such an oscillator has been exapted to various modalities, behaviors and way of moving.

## Introduction

Navigating through space implies both to acquire information (exploration) and to use this information to move in the correct direction (exploitation). A way to acquire information is to sample the environment by actively moving. Such ‘active-sampling’ is common across the animal kingdom spanning from invertebrates (Dittmar et al., 2010; Gomez-Marin et al., 2011; Lehrer, 1996) to vertebrates (Dawkins and Woodington, 2000; Otero-Millan et al., 2008; Wachowiak, 2011; Wesson et al., 2008; Wolf et al., 2017). However, active sampling entails movements that are typically different than moving toward the goal and thus require the animal to solve a tradeoff. Achieving a balance between sampling actions (exploration) and goal-directed actions (exploitation) lies at the core of the control of behavior but how it is achieved and evolved in animals remains unclear.

One way to solve the exploration/exploitation tradeoff is to move in order to maximize the information gain, for instance by using the so-called infotaxis (Hernandez-Reyes et al., 2021; Vergassola et al., 2007). Climbing up an information gradient can lead directly towards the goal if the source of information is emitted by the goal itself, such as when targeting sound or odor sources. However, it is unclear how such a strategy could apply when one moves without perceiving its final goal, such as during visual route following.

Another way of sampling the world is through the intrinsic production of regular alternations between left and right turns along the path: lateral oscillations. Lateral oscillations are observed in wide range of taxa (vertebrates: DeBose and Nevitt, 2008; Wolf et al., 2017; invertebrates: Freas and Cheng, 2022; Iwano et al., 2010; Kanzaki et al., 1992; Namiki and Kanzaki, 2016; Wystrach et al., 2016).

Oscillatory paths have been mainly studied in the context of olfaction, whether during plume following in moths (Kuenen and Baker, 1983; Olberg, 1983; Kanzaki et al., 1992; Kuenen and Baker, 1983; Namiki et al., 2014), trail following in ants (Hangartner, 1967) or odor gradient climbing in *Drosophila* larvae (Wystrach et al., 2016) and *Caenorhabditis elegans* (Izquierdo and Lockery, 2010). The intrinsic production of oscillations enables an efficient sampling of odor across locations and models show that oscillations modulated by odor perception enables to reach the source in a remarkably effective way (Adden et al., 2020; Izquierdo and Lockery, 2010; Wystrach et al., 2016).

Beyond that, modeling also shows that the production of oscillations can be equally effective during visual navigation tasks. For instance, having the amplitude of oscillation modulated by familiarity of visual scenes can produce robust visual navigation behavior such as route-following (Kodzhabashev and Mangan, 2015) or homing (Le Möel and Wystrach, 2020). It also captures particular behavioral signatures observed in ants (Murray et al., 2020). Moreover, visually guided insects such as ants (Graham and Collett, 2002; Lent et al., 2013, 2010; Jayatilaka et al., 2017; Murray et al., 2020), wasps (Stürzl et al., 2016; Voss and Zeil, 1998) or bumblebees (Philippides et al., 2013) do display oscillations, whose expression appears to be coupled with visual perceptual cues. But whether these lateral oscillations are produced internally, how exactly they interact with visual cues, and how they participate in the exploration/exploitation tradeoff during a visual navigational task remains unclear.

Here, we focused on the expression of oscillations in two ecologically and phylogenetically distant ant species known to use vision for navigation (*Myrmecia croslandi* and *Iridomyrmex purpureus*). These insects are known to rely on two mains strategies to guide their foraging journeys to an inconspicuous goal (Nardendra et al., 2013; Card et al., 2016, see also SI1). The first strategy, commonly called path integration (PI), allows individuals to continuously estimate the distance and compass direction that separates them from their nest (or other starting points) during their foraging trip (Collett and Collett, 2017; Heinze et al., 2018; Muller and Wehner, 1988). The second, commonly called view-based navigation, involves the learning and subsequent recognition of learnt visual panorama (Collett and Cartwright, 1983; Wehner, 1979; Zeil, 2012). To see how these navigational strategies do interact with oscillation, and how they participate to the exploration /exploitation tradeoff during a visual navigational task, we mounted ants of both species on a trackball device (Dahmen et al., 2017). This device enabled us to record their motor behavior and control the visual cue perceived directly in their natural environment. We characterized in detail whether the obtained trajectories show a regular oscillatory pattern of movements and how these patterns are influenced by the presence or absence of: (1) visual input, (2) learnt visual terrestrial cues, (3) path integration homing vector and (4) rotational visual feedback. In both species our results revealed the presence of a conserved pattern of oscillations generated intrinsically, which comprises both angular and forward velocity components. The amplitude of this dynamic signature is modulated by visual information in a way that is adapted to the navigational tasks at hand. Finally, a simple neural circuit model of reciprocal inhibition between left and right pre-motor area can readily explain the production of these movement dynamics.

## Results

### Ants mounted on the trackball display their natural navigational behavior

We first investigated whether ants mounted on the trackball display the expected, natural navigational behavior for different experimental conditions. When released on the ground in their natural environment, both species studied here (*Myrmecia croslandi* and *Iridomyrmex purpureus)* are known to rely on learnt terrestrial cues as well as, to a lesser extent, on PI (Card et al., 2016 see Fig. S1 for preliminary experiment on *I. purpureus*; Murray et al., 2020; Narendra et al., 2013; Zeil et al., 2014). Here we mounted ants mounted on the trackball in a way that enabled them to physically rotate and control their actual body orientation (Dahmen et al., 2017). The trackball was placed in three distinct visual conditions: (1) along the individual ant’s familiar route (F) therefore in presence of familiar terrestrial cues; (2) in an unfamiliar location (U) 50 meters away from their usual route; or (3) without any visual input: in complete darkness (D). For each of these visual conditions, ants were tested either with (full-vector (FV) ants) or without (zero-vector (ZV) ants) path integration information. While FV ants were captured at their feeding place (from their foraging tree (*M. croslandi*) or from their trained feeder (*I. purpureus*)) and thus possess a PI homing vector pointing in the food-to-nest compass direction, ZV ants were caught just before entering their nest and thus no longer possess a PI homing vector.

For both species and irrespectively of the PI state (FV or ZV), ants mounted on the trackball within their familiar visual route (i.e., in the presence of learnt terrestrial cues) displayed paths that were oriented toward the nest direction (Fig. 1A, B first column; Rayleigh test with nest as theoretical direction p*s*: *M. croslandi*: FV & ZV < 0.001; *I. purpureus*: FV & ZV < 0.001), proving that they recognized and used the familiar terrestrial cues to orient. When tested in unfamiliar surroundings, FV ants of both species were oriented toward the theoretical direction of the nest as indicated by their PI (Fig. 1A, B second column, first row; Rayleigh test with PI as theoretical direction p*s*: *M. croslandi*: FV< 0.001; *I. purpureus*: FV< 0.001); but ZV ants showed random orientations (Rayleigh test p*s*: *M. croslandi*: ZV=0.443; *I. purpureus*: ZV=0.672). This confirms that the distant release points were indeed unfamiliar to the ants and that both species relied on their PI when tested as FV ants. When tested in total darkness, ants displayed randomly orientated paths in all conditions (Fig. 1A, B third column; Rayleigh test p*s*: *M. croslandi*: FV=0.354; ZV=0.360; *I. purpureus*: FV=0.213, ZV=0.866), showing that the chosen dark condition was effective in preventing ants using terrestrial or celestial cues for orientation. Overall, these results demonstrate that both species use learnt terrestrial cues as well as their PI to navigate when mounted on the trackball system (Fig. 1A, B).

**Figure 1.**
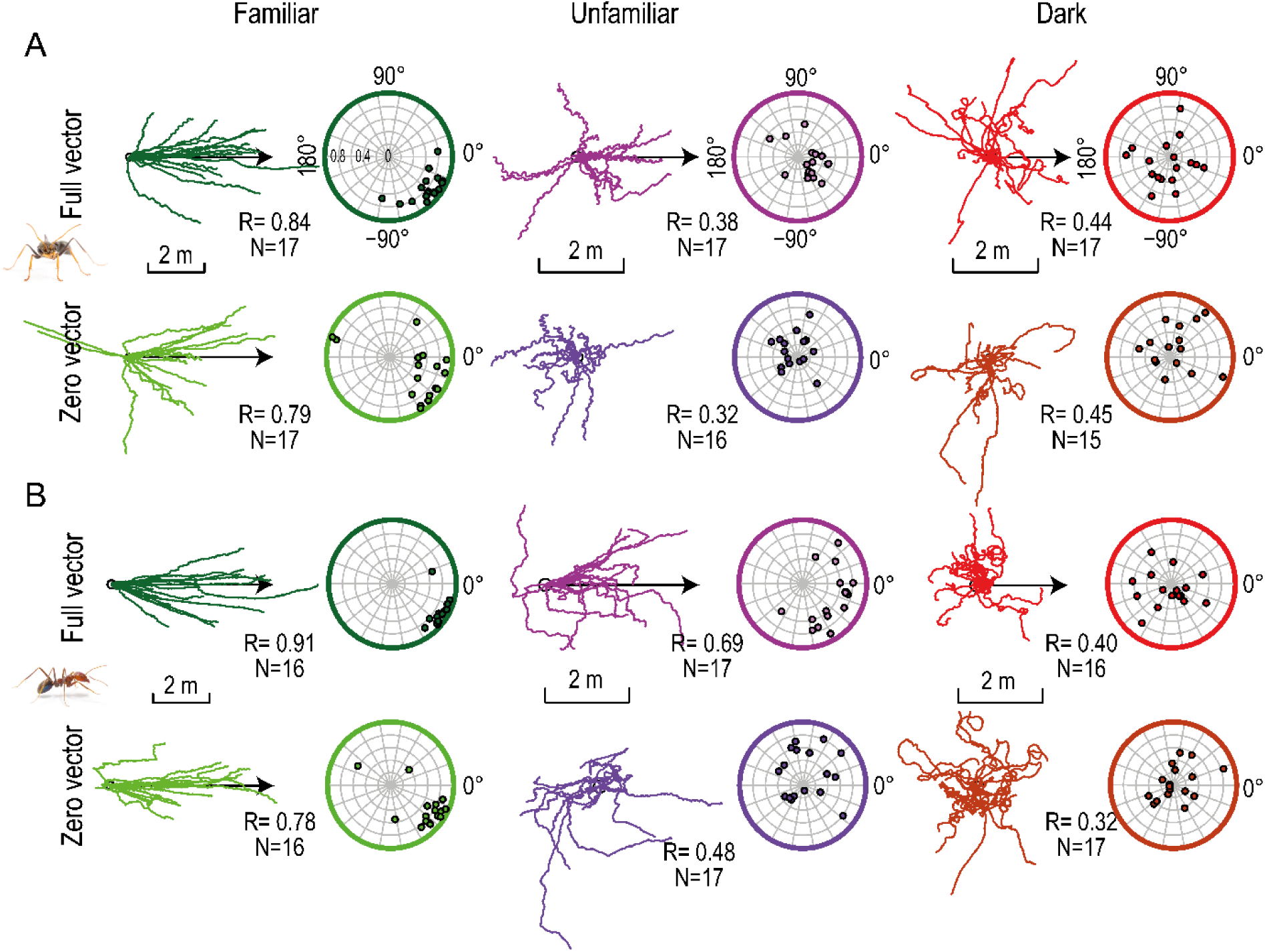
Ants on the trackball display natural navigational behaviors. Data presented for both *M. croslandi* (A) and *I. purpureus* (B), tested in a familiar (green) or an unfamiliar panorama (purple) and in the dark (red). For each condition, individuals were tested either with (full vector) or without (zero vector) path integration information. The first column of each condition shows the paths oriented to the theoretical direction of the nest (arrow). In the second column, each dot indicates the average circular vector calculated over the entire path of an individual, showing the mean direction and the average vector length (i.e., a point closer to the periphery indicates straighter paths). The R values indicate the length of the average resulting vector of the population. The direction (in familiar terrain) or theoretical direction (for FV in unfamiliar terrain) of the nest is 0°. Ants photography credit: Ajay Narendra.

Independent of the PI state, the ants’ travel directions were more constant in familiar terrain as compared to the other conditions (R values in Fig. 1A, B; Familiar vs. Unfamiliar & Familiar vs. Dark, both species: p*s* < 0.001), showing that the recognition of learnt terrestrial cues was most effective to maintain a constant direction of travel. With regards to the effect of PI, FV ants tended to keep their direction more constant than ZV ants but this effect was small, and reached significance only in unfamiliar terrain for *I. purpureus* (Fig. 1; FV vs. ZV p = 0.049) but not in familiar terrain (Fig. 1; FV vs. ZV both species: p*s*≥0.0589). Finally, there was no difference between the PI state for both species in the dark condition (Fig. 1; FV vs. ZV p*s*=1). Therefore, both species do use their PI to orient in unfamiliar terrain, but it seems that PI has a limited impact on their path’s straightness.

These results are consistent with what is observed in these species when naturally walking on the floor (Card et al., 2016; Narendra et al., 2013).

### Ants display regular lateral oscillations

To determine whether ants display regular lateral oscillations – that is, alternate between left and right turns at a steady rhythm – we first used our video recordings to track the ant’s change in body orientation when mounted on the trackball. The body angular velocity signal is independent of the ant’s actual forward movement and thus a direct reading of its motor control for turning. We conducted a Fourier analysis on autocorrelation coefficients of the angular velocity time series to obtain a ‘power spectral density’ (see Fig. S2 for detailed method). The obtained Fourier’s magnitudes are a measure of the regularity of the rhythms in the time series, with high magnitude values indicating regular oscillation at the corresponding frequency (Fig. S2). We thus selected the frequency corresponding to the highest magnitude (peak), that is, the most salient rhythm in the signal. In all experimental conditions, the obtained peak fell in the expected range of 0.2 to 1.5 Hz, as observed in other insect species (Lonnendonker, 1991; Wystrach et al., 2016). It should be noted that this rhythm is 10 to 50 times slower than ants’ typical stepping frequency (Zollikofer, 1994) and thus are not the consequence of their rhythmic walking gait per se but the product of an additional oscillatory mechanism. The obtained Fourier magnitudes were higher than those obtained with the spectral density of a Gaussian white noise (Fig. 2A, B dashed line; Wilcoxon one-tail test: p*s*≤2.144e-03), showing that ants displayed lateral oscillations with a higher regularity than random.

**Figure 2.**
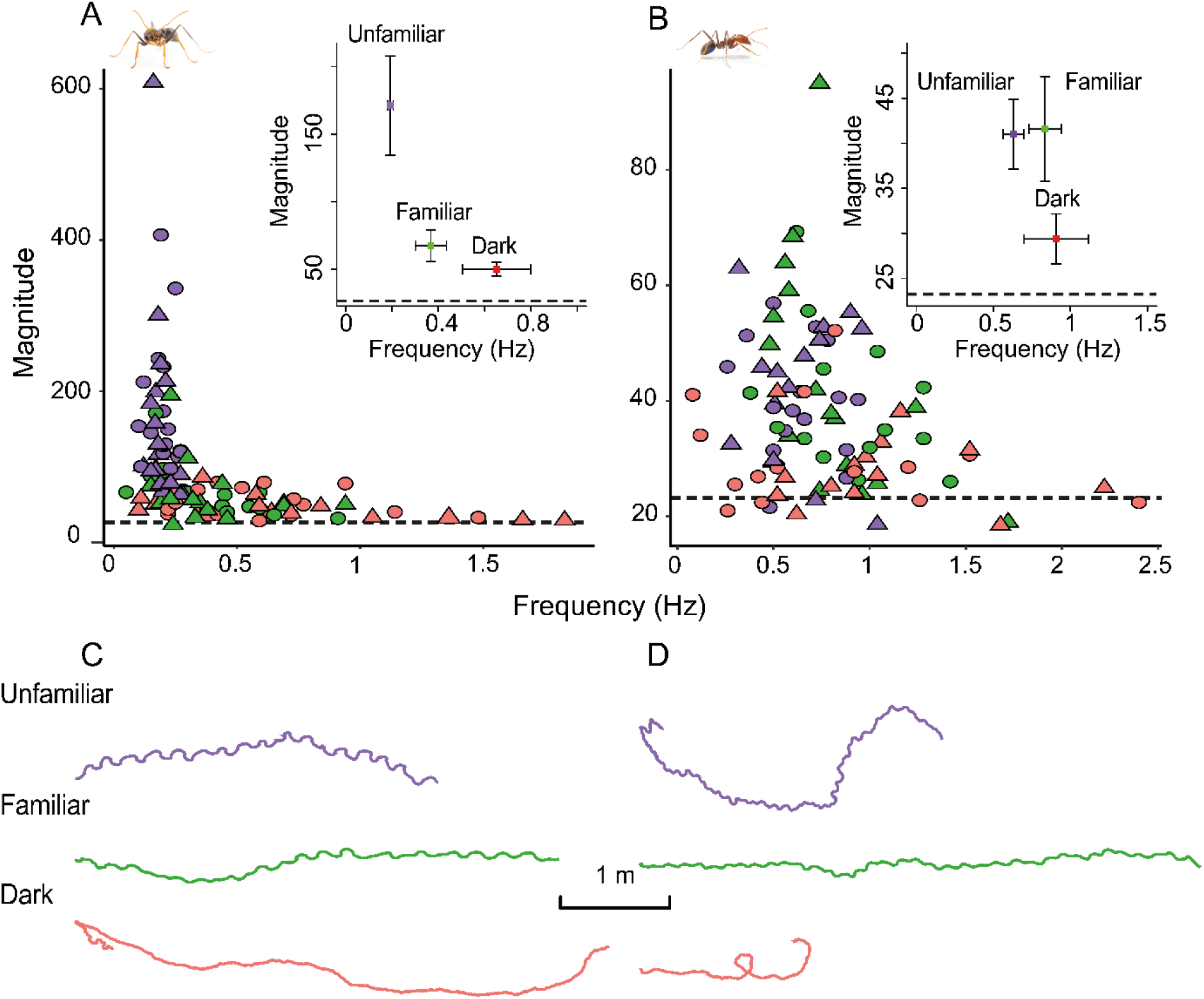
Oscillation characteristics vary across visual conditions. The graphs show the frequency and spectral density magnitude of the dominant oscillation (highest magnitude) in *M. croslandi* (A) and *I. purpureus* (B) (for method see method see Fig. S2 A, C, D, E). High frequencies indicate a fast-oscillatory rhythm and high magnitudes indicate a strong presence of this oscillation. Individuals were tested within a familiar panorama (green), an unfamiliar panorama (purple) or in the dark (red). Symbols indicate whether the state of the path integration vector was full (round) or set to zero (triangles). Inserts show the mean frequency against the mean magnitude of each visual condition with the associated 95% confidence interval around the mean. The dashed black line represents the mean of the spectral density peak magnitudes resulting from 200 Gaussian white noise signals. The second row show an example path for each condition for M. *croslandi* across 100 s (c) and *I. purpureus* across 83 s (D). Ants photography credit: Ajay Narendra.

### The navigational context modifies the regularity of oscillations

We analyzed whether our different test conditions influenced the magnitude of the individuals’ spectral density peaks, which indicates how regular the oscillations are. The statistical model revealed no interaction between the effects of the visual panorama type and the state of the path integrator (*M. croslandi*: AIC=-198.8, F_2,101_=0.434, p=0.647; *I. purpureus*: AIC=-167, F_2,90_=0.232, p>0.79). The additive effect model (i.e., without interaction, *M. croslandi*: AIC=-215.8; *I. purpureus*: AIC=-170.5) explains the variation of the magnitude peaks relatively well (R^2^ *for M. croslandi*: marginal=56% & conditional=67%; *I. purpureus*: 16%). The subsequent Anova revealed that the visual panorama type has a significant effect on the regularity of oscillations in both species (panorama effect for *M. croslandi*: F_2,101_=83.663, p < 0.001; *I. purpureus*: F_2,90_=10.038, p < 0.001); however, there was no overall effect of the PI state (vector effect: *M. croslandi*: F_1,101_=1.291, p>0.200; *I. purpureus*: F_1,90_=0.012, p>0.900).

For *M. croslandi*, the post-hoc analysis revealed that the magnitude peak varied significantly across the three visual conditions (Fig. 2A, F vs. U: p<0.001; F vs. D: p=0.016; U vs. D: p<0.001). These differences in peaks magnitudes show that the oscillations are most regular in unfamiliar terrain, intermediate in familiar environment and least regular in the dark (mean±se: U_(FV+ZV)_=171±19; F_(FV+ZV)_=68±6; D_(FV+ZV)_=50±3). Interestingly, the effect of individuality is significant in *M. croslandi* (p=0.015), indicating that some individuals have a better consistency in their oscillations than others, across conditions.

For *I. pupureus*, the post hoc analysis reveals that the regularity of the oscillations does not differ between familiar and unfamiliar terrain (Fig. 2B, p>0.96, df=86, mean±se: F_(FV+ZV)_ =41.59± 3; U_(FV+ZV)_=41±2) but both yield more regular oscillations than in the dark (p< 0.001, mean±se: D_(FV+ZV)_=29.4±×1.45).

Overall, these differences confirm the existence of regular oscillations in both unfamiliar and familiar terrain, regardless of the PI state. In the dark however, peak magnitudes are significantly weaker, closer to Gaussian noise and with a wider frequency range, suggesting that the Fourier’s peak in some individuals may result from noise (Fig. 2A, B). If oscillations persist in the dark, their expression is at least greatly inhibited (Fig.2).

### The navigational context modulates the oscillation frequency

*Myrmecia* foragers tested in familiar panorama showed no differences in oscillation frequencies across PI states (Wilcoxon test for repeated measures; F: FV vs. ZV p=0.635; U: FV vs. ZV p=0.343). Again, independent of the PI state, the oscillatory frequencies were significantly higher in familiar than in an unfamiliar visual panorama (p=0.023, mean±se: F_(FV+ZV)_ =0.37±0.036 Hz; U_(FV+ZV)_ =0.19±0.008 Hz; Fig. 2A, C). *Iridomyrmex purpureus* showed the same tendency to oscillate quicker in familiar than in unfamiliar terrain (mean±se: F_(FV+ZV)_ =0.83±0.056 Hz; U_(FV+ZV)_ =0.63±0.037 Hz; Fig. 2B, D), but this effect did not reach significance (p*s* ≥ 0.180).

Thus, the presence of familiar visual cues tends to increase the frequency of oscillations, at least for *M. croslandi* (Fig. 2A, C), but the state of the PI vector (FV or ZV) had no observable effect (mean±se in F: FV=0.38±0.05 Hz, ZV=0.34±0.04 Hz; U: FV=0.19±0.01Hz, ZV=0.18±0.0009 Hz).

### The navigational context modulates the angular and forward velocity of oscillations

To investigate the dynamics of the ant oscillatory movements in more detail, we combined for each ant its body angular velocity signal – obtained from video recording – with the forward movement signal – obtained from the trackball movements. We reconstructed ‘average cycles’ for each individual by pooling the angular and forward velocity recordings 3s before and 3s after the moments where ants switch from a left to a right turn (when the time series of the angular velocity crosses zero from negative to positive; see Fig. 3—figure supplements 1). That way, we can quantify each individual’s average dynamic of angular and forward velocity and compare them across conditions.

For *M. croslandi*, the visual surrounding had a strong effect (p<0.001): ants displayed higher angular velocities (Fig. 3A, inset boxplot upper row; mean±se: U_(FV+ZV)_=129±4.8 deg/s; F_(FV+ZV)_=67.5±4.4 deg/s) and higher forward velocities (Fig. 3A, inset boxplot second row, mean±se: U_(FV+ZV)_=4.8±0.34 cm/s; F_(FV+ZV)_=2.3± 0.2 cm/s) in an unfamiliar environment. There was however neither an effect of the PI state on turn or forward velocity (p>0.05) nor an interaction between the visual surrounding and the PI state (mean peak of angular velocity model: AIC=160.9, interaction p>0.05; amplitude of forward velocity model: AIC=109.9, interaction p=0.22). Therefore, the amplitude of lateral oscillation within the path are higher in an unfamiliar panorama than in a familiar one independently of the PI sate (Fig. 2 C). Interestingly, *M. croslandi* ants showed strong individual idiosyncrasies across conditions (forward velocity peak amplitude: random effect, p<0.05, R2 marginal=0.39, R2 conditional=0.61; angular velocity peak amplitude: random effect, p>0.05, R2 marginal=0.5; R2 conditional=0.66).

**Figure 3:**
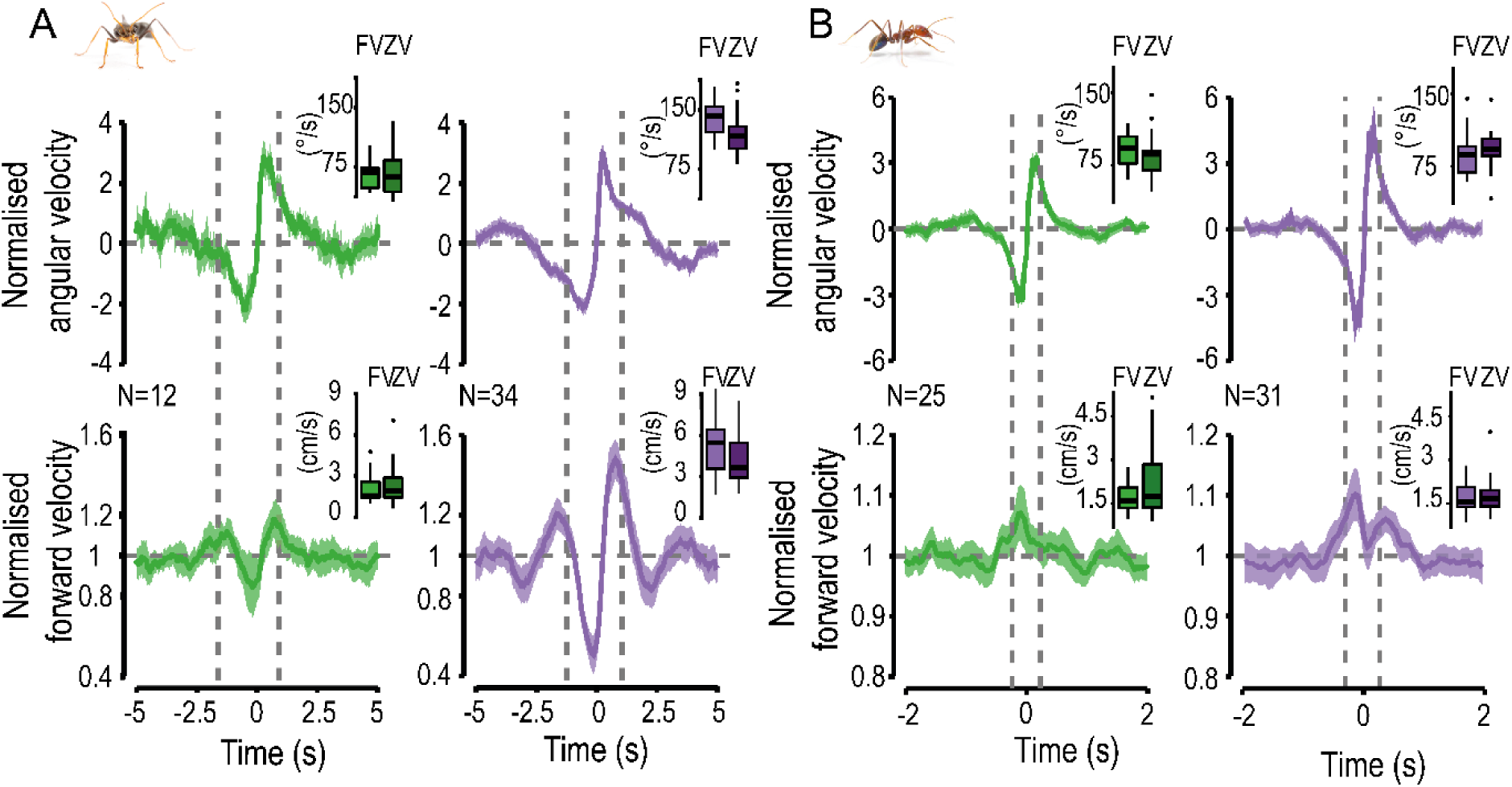
Population-average cycle of oscillations shows how angular and forward velocities co-vary. Angular velocities (top-row) and forward velocity (bottom row) co-vary in a way that seem conserved across species (*M. croslandi* (A) and *I. purpureus* (B)) when on familiar route (green) or in unfamiliar terrain (purple). Population cycles have been reconstructed by merging full-(FV) and zero vector (ZV) data and normalizing the data amplitude within the individual’s average cycle (see Fig. 3 —figure supplements 1). Color areas around the mean curves represent the 95% confidence interval, based on the inter-individual variation. Dashed lines represent the moment when the ants are facing their overall direction of travels. Insert boxplots show the actual distribution of the non-normalized mean peak of angular speed and forward velocity amplitude for the individuals’ average cycle (FV on the left, ZV on the right). Ants photography credit: Ajay Narendra.

**Figure 3—figure supplements 1:**
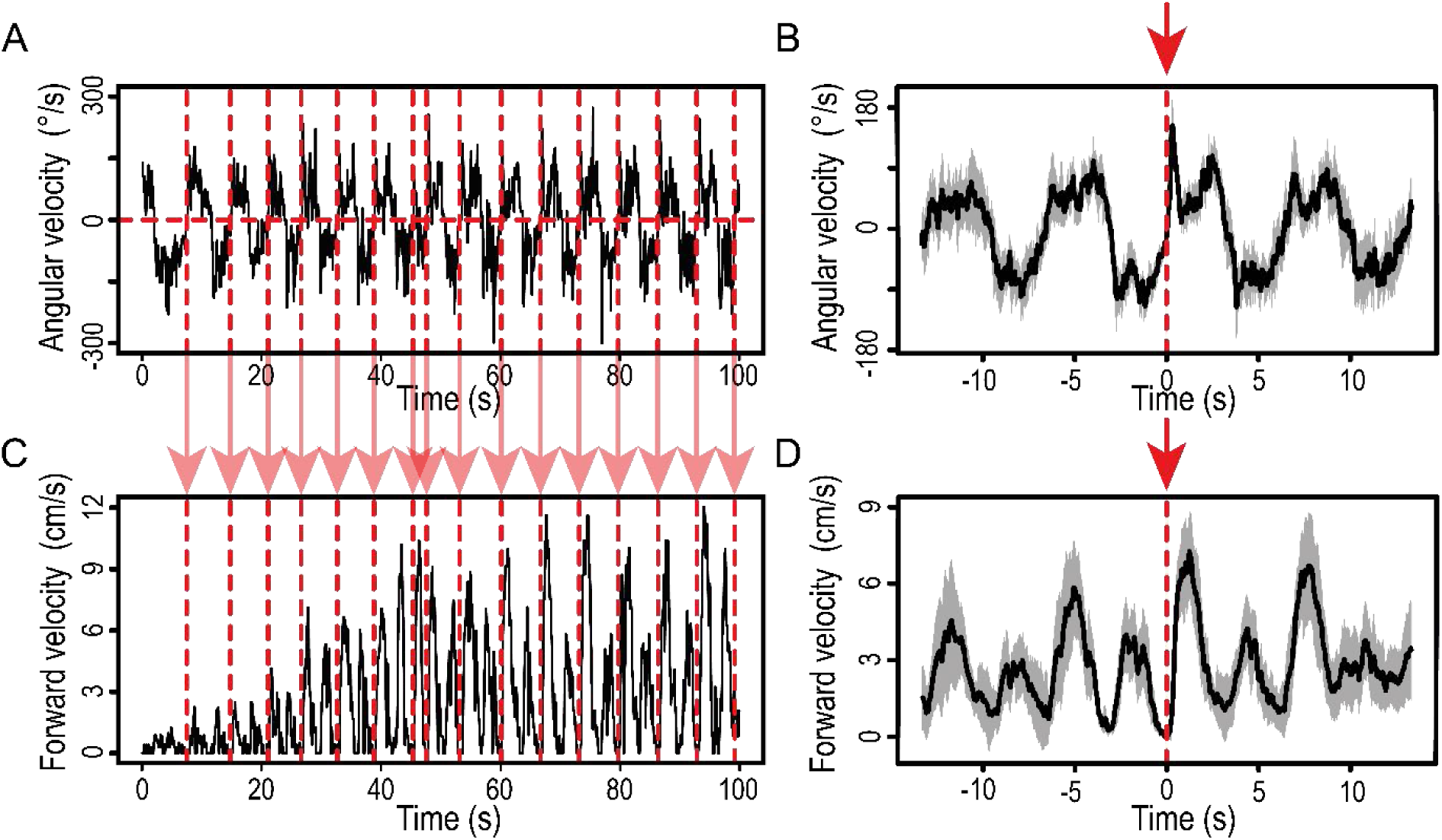
Methodology to obtain an average oscillation cycle. (A) We used the angular velocity signal to flag moments when the signal crosses 0 and goes up to positive angles (vertical dashed red lines). (B) We extracted portions of the signal of ±3s (large enough to contain a full oscillation cycle) around each of the flags, aligned them at flag = t_0_ and averaged them to obtain the individual’s average cycle (±95% confidence interval in grey) of the angular velocity signal. (C, D) The individual’s average cycle for the forward velocity signal was achieved using the angular velocity flags (vertical dashed red lines as in A).

For *I. purpureus* ants, none of the models reached significance (p*s* >0.05); showing that the dynamics of angular velocities and forward velocity underlying the oscillations are rather constant across navigational context (Fig. 3B insert). Showing no clear effect on the amplitude of oscillation display withing the path (Fig. 2D).

### The conserved dynamics of lateral oscillations

To test for the existence of conserved oscillatory dynamics across individuals, each individual average cycle (Fig. *3*—figure supplements 1 for method example) was normalized and averaged to obtain a cycle at the population level. To ensure that we pooled ants oscillating at similar frequency ranges, we separated data from familiar and unfamiliar terrain and selected only individuals showing a peak frequency within 0.1 to 0.27 Hz for *M. croslandi* (N=12) and within 0.26 to 1.04 Hz for *I. purpureus* (N=25). These frequency ranges correspond to high magnitude, ensuring that they are not a consequence of noise. The emerging population-averaged oscillation patterns show the existence of movement dynamics that are consistent across individuals of both species, for both familiar and unfamiliar environments (Fig. 3, Fig. 4). Forward velocity co-varies with angular velocity in a particular way. Forward velocity is quite low when the ants reverse their turning direction (i.e., when angular velocity crosses zero). In contrast, we observe two significant peaks of forward velocity, which happen briefly (up to 1s) while the ant is sweeping to the left or right. Remarkably, these peaks coincide well with the moment when the ants’ body orientation is aligned with its overall direction of travel (Fig. 3, dashed lines) even though during these moments the ants’ angular velocity is quite high. In other words, ants display large sweeps from one side to the other and do not maximize the time they spent facing their goal direction, which could be achieved by reducing angular speed at this moment. However, they increase their forward velocity precisely while doing so.

**Figure 4:**
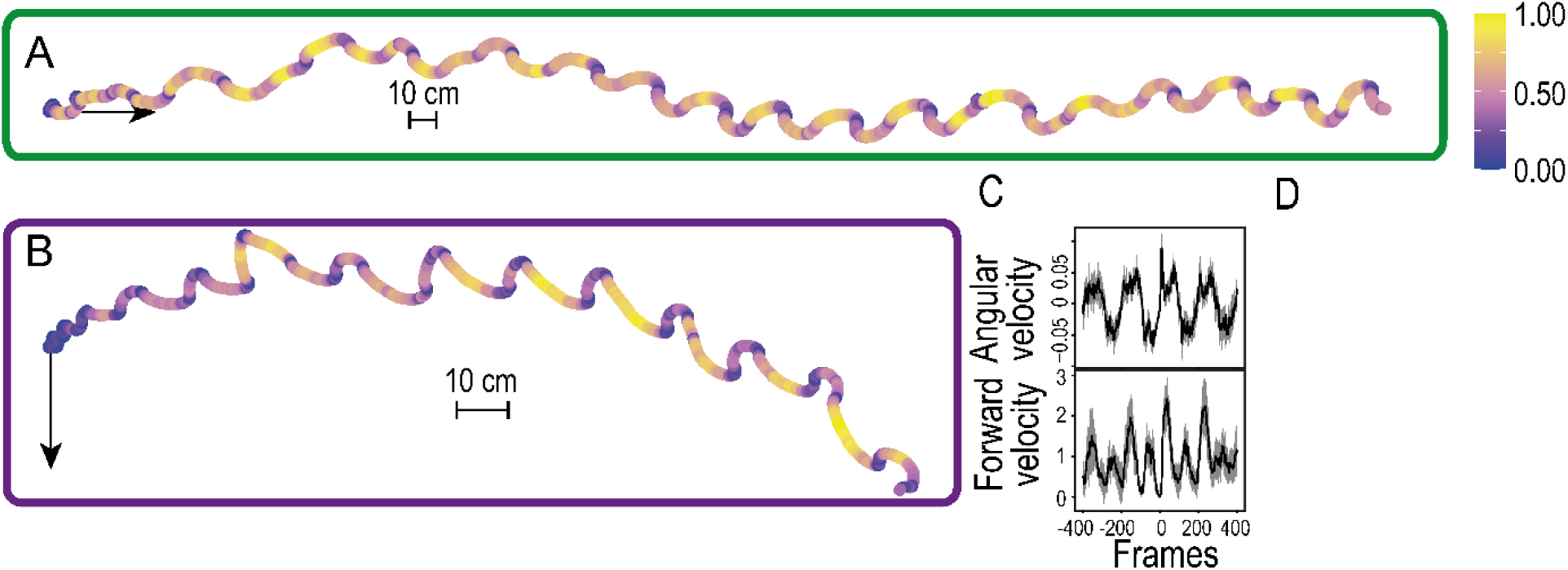
Dynamics of forward velocity are the same in familiar and unfamiliar panoramas. Reconstructed path (100 s) of *M. croslandi* individuals recorded in a familiar (A) and in an unfamiliar (B) panorama. Each path has been colored according to the forward velocity data (yellow forward movement; blue no forward movement). On both paths, the arrow indicates the theoretical direction of the nest. (C) Averaged cycle of the individual recordings in B, showing that forward and angular velocities’ peaks happen simultaneously. (D) Video of the individual corresponding to the paths shownin B.

To ensure that these dynamics are not a consequence of being mounted on the trackball set-up, we replicated the same analysis using tracks of *M. croslandi* ants recorded directly on the ground while displaying learning walks around their nest (courtesy of Jochen Zeil, see Zeil and Fleishmann 2019). The natural learning walks also showed regular oscillations as well as the same dynamical relationship between angular and forward velocities (Fig. 5). Ants walking on the ground also tend to increase their forward velocity when they are aligned with their general direction of travel.

**Figure 5.**
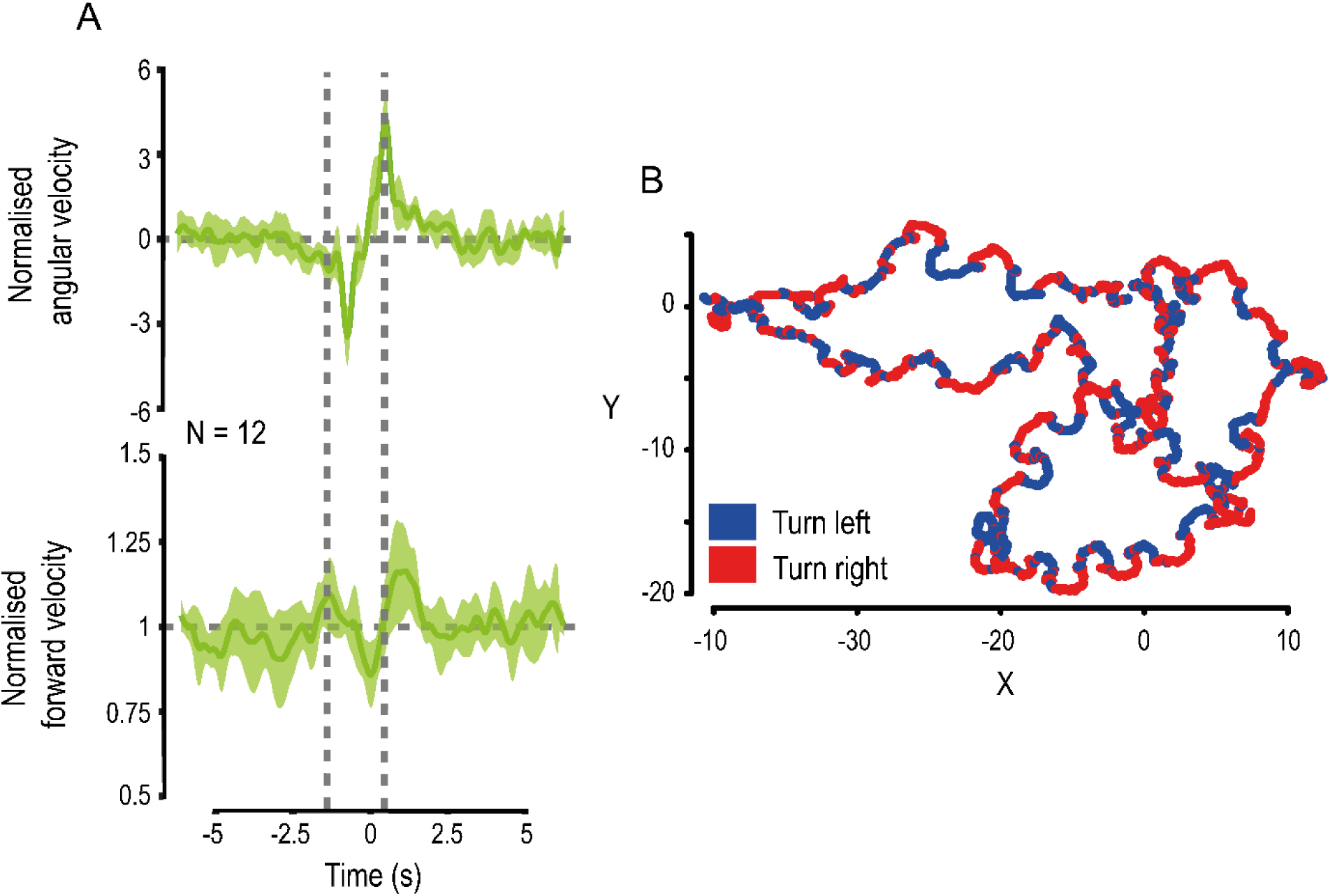
Ant learning walks show similar dynamics of oscillation. Ants path and head orientation were recorded (25 Hz) during their learning walks directly on the ground. (A) Angular velocity (top-row) and forward velocity (bottom row) co-vary in a similar way than when recorded on the track ball (see Fig. 3). Population cycles have been normalized by the data amplitude within the individuals mean cycle (see Fig. *3*—figure supplements 1 for example). Color areas around the mean curves represent the 95% confidence interval, based on the inter-individual variation. (B) Example path of an individual showing clear alternation between right (red) and left (blue) turn.

### Oscillations are modulated by rotational feedback

To test whether oscillations result from an intrinsic oscillator rather than a servo-mechanism based on sensory information, we investigated whether ants deprived of rotational feedback (either via optic-flow or compass cues) would still oscillate. For that, ants were mounted on the trackball in a way that prevented them from physically rotating their body on the ball: when the ant tries to turn, it is the ball that counterturns. In other words, the fixed ant experiences neither a change of body orientation nor visual rotational feedback when trying to turn (see Fig. S2B). For this experiment, we used *M. croslandi* ants in unfamiliar terrain as this condition produced the clearest oscillations in our previous experiments (Fig. 2). To control for potential effects of facing directions (depending on celestial or terrestrial visual cues), we tested each ant fixed in eight subsequent different orientations with a 45° rotational shift.

Despite the absence of visual rotational feedback, the obtained peak magnitudes of the angular velocity spectral density (Fourier analysis on autocorrelation coefficient, see Methods) were much higher than expected from a Gaussian white noise (p<0.001) showing clearly the presence of a regular alternation between left and right turns (Fig. 6). The ants’ body orientation relative to the world had no effect on the magnitudes (orientation: F_7,83_=0.3729, p=0.914) nor did the other random parameters (individual p=0.933, sequence p=0.473). The mean oscillation frequencies (mean±se: 0.17±0.007 Hz) were quite close to what we observed when these ants were free to rotate in an unfamiliar environment (Wilcoxon test for repeated measures: p*s*=1, N=88, Fig. 6). Thus, ants can display regular turning oscillations without rotational feedback, whether as a result of optic flow or a change of orientation relative to directional cues such as the visual panorama, wind, celestial compass cues or the magnetic field. Consequently, we are left with the conclusion that these oscillations are generated intrinsically.

**Figure 6.**
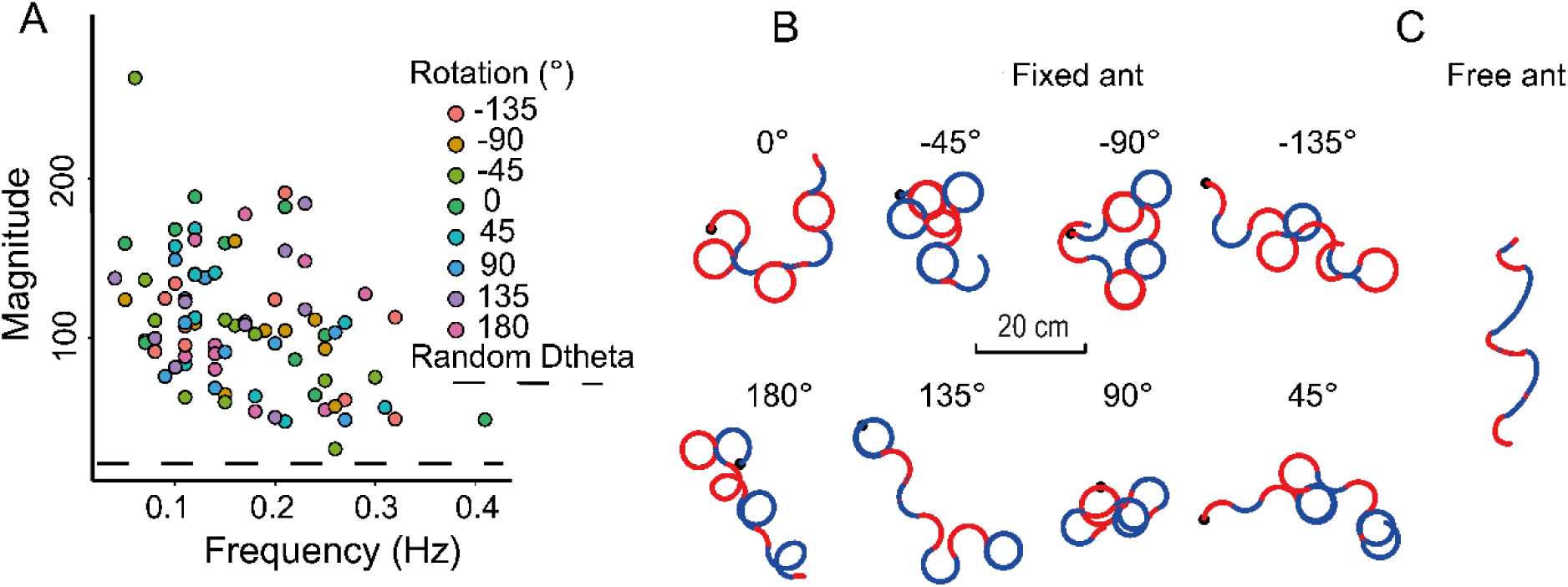
Ants still oscillate in the absence of rotational feedback. Zero vector ants were tethered on the trackball in a way that prevented actual body rotation; and teseted in an unfamiliar environment. (A) Distribution of the individual Fourier dominant peak frequency and magnitude based on the angular velocity’s spectral density time series (as in Fig. 2, method see Fig. S2 B, C, D, E). High frequency indicates a fast-oscillatory rhythm, and a high magnitude indicates a strong presence of this oscillatory rhythm. The color symbols represent the rotation relative to the theoretical nest direction (0°). The dashed black line represents the mean magnitude obtained from 200 Gaussian white noise signals (18.75). (B) Example paths of different individuals fixed in different orientations. (C) Example path from an ant that was free to rotate on the trackball, recorded in unfamiliar terrain in previous experiments. (B, C) In both situations, ants alternate regularly between right (blue) and left (red), but turns are sharper in the absence of rotational feedback.

Interestingly, the absence of rotational feedback led to higher angular velocities (mean: fixed=176 deg/s; free=128 deg/s; p<0.001), higher forward velocity (mean: fixed=11 cm/s; free=5 cm/s; p<0.001) and slightly slower turn alternation (i.e., lower frequency: fixed=0.16 Hz; free=0.19 Hz, p=0.0189) than ants that were free to physically rotate on the ball (Fig. 7). Rotational sensory feedback is thus involved in limiting the amplitude of the oscillations.

**Figures 7:**
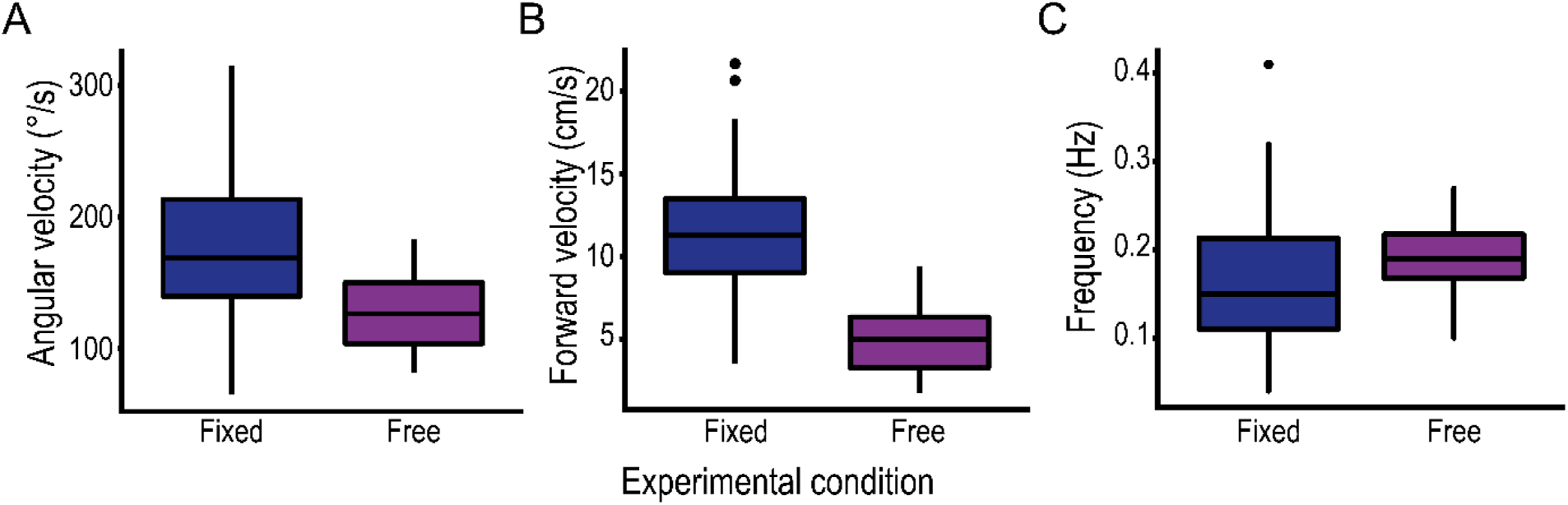
Absence of visual rotational feedback increases the amplitude of oscillation. Distribution of *M. croslandi* individuals’ average oscillation dynamics in unfamiliar terrain when recorded on the track ball either in fixed mode preventing rotational feedback (blue) or free (purple). Peak of angular velocity (A), peak of forward velocity (B) and frequency (C) of the individual’s average oscillation cycle show that ants move faster, turn faster and slightly longer in the absence of rotation feedback.

### The observed movement signature emerges readily from a simple neural circuit

We designed a simplistic neural model based on the circuitry of the insect pre-motor area so called lateral accessory lobes (LAL) (Namiki and Kanzaki, 2016a; Steinbeck et al., 2020). The purpose of this model is not to match observed data quantitatively (Adden et al., 2020) but simply to test whether the co-varying relationship observed between angular and forward velocity can readily emerge from this type of circuit. Activation of the neurons in the left and right pre-motor areas are known to mediate left and right turns respectively in insects (Berni, 2015; Berni et al., 2012; Iwano et al., 2010; Kanzaki, 2005; Namiki and Kanzaki, 2016b; Steinbeck et al., 2020). Furthermore, these left and right regions are known to reciprocally inhibit each other (Iwano et al., 2010; Berni et al., 2012; Namiki and Kanzaki, 2016b; Steinbeck et al., 2020). Modelling two reciprocally inhibiting neurons with internal feedback –that tries to maintain a basal activity (Fig. 8)– is sufficient to obtain the typical oscillatory activity between left and right LAL (Iwano et al., 2010; Steinbeck et al., 2020), and thus provides an explanation for the regular oscillations between left and right turns observed in insects. Interestingly, we show here that simply assuming that forward velocity is controlled by the sum of left and right output – while angular velocity results from their difference (Adden et al., 2020; Steinbeck et al., 2020; Wystrach et al., 2016)– is sufficient for the observed covariation to emerge (Fig. 8). The agent accelerates while facing its general direction of travel and slows down at the end of its sweeping movement when its orientation extends most to the left and the right (Fig. 8C, 4). What’s more, the emergence of this particular relationship is robust to parameter change (Figure 8—figure supplements 1) and thus a stable feature of either these types of circuits.

**Figure 8.**
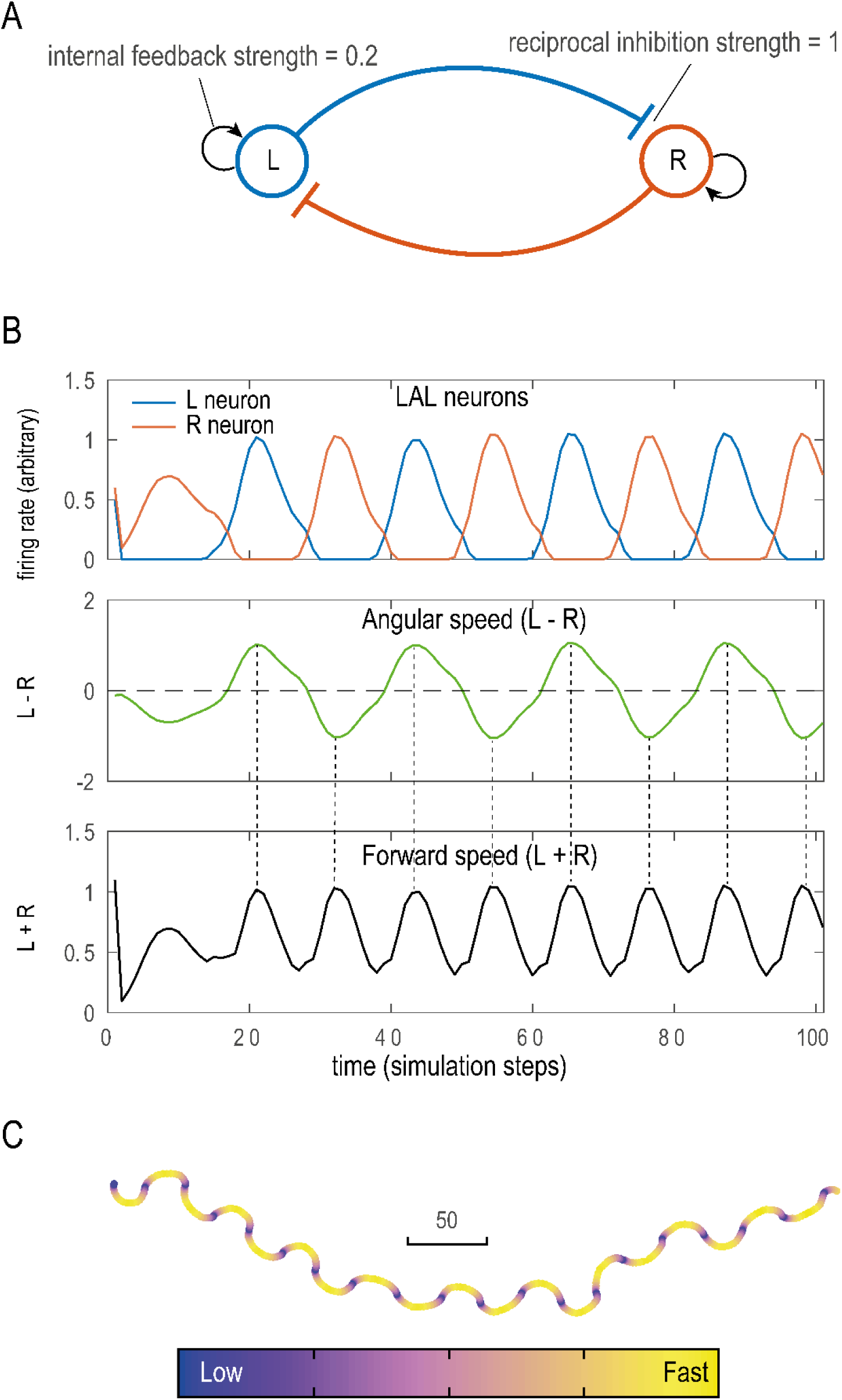
Reciprocal inhibition between two units produces the observed relationship between angular and forward velocity. (A) Scheme of the model abstracted from the lateral accessory lobe (LAL). The Right (R) and Left (L) hemisphere neurons reciprocally inhibit each other (blue and red connection), while trying to sustain a basal firing rate through internal feedback (black arrow). (B) This results in the emergence of stable anti-phasic oscillatory activity between the R and L neurons (B, upper row). Assuming that angular velocity is controlled by the difference (B, middle row) – and forward speed by the sum (B, bottom row) – between left and right activation is sufficient to elicit the movement dynamics observed in ants (see Fig. 3C). As ants, the model results in a zigzagging path where velocity drops during turn reversal, and increases when facing the overall direction of travel (Fig. 3, 4). While the model enables the modulation of amplitude and frequency of the oscillations, the forward/angular velocities relationship is quite robust to parameter change (Fig.8—figure supplements 1).

**Figure 8—figure supplements 1.**
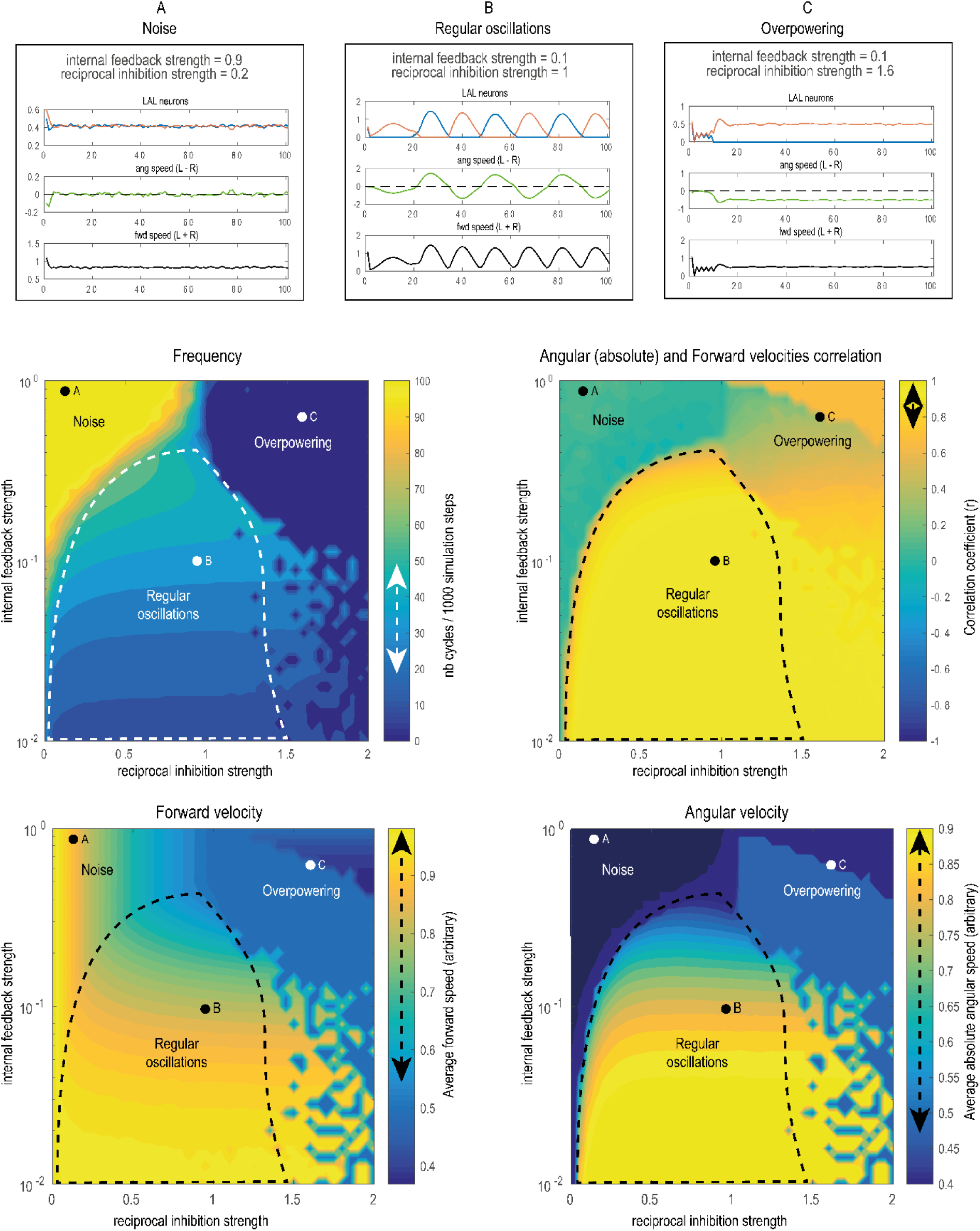
The covariation between angular and forward velocity is robust to parameter change. Exploration of our abstract LAL model’s (figure 8) sensitivity to change in the two free parameters (inhibition strength and internal feedback strength). Regular oscillations between LAL neurons emerge in a wide space of parameters (functional range delineated by the dashed line, an example for a given parameter set (dot) is shown in (B)). While frequencies (right of middle row), Forward and Angular velocities (bottom row) can vary roughly across a factor 2 across the functional parameter range, the correlation between absolute angular and forward velocity is constrained around 1 (dashed arrows in the color bars indicate the variation in the functional range). This positive covariation indicates that the agent slows down (low forward velocity) when reversing turning direction (low absolute angular velocity) and accelerates (high forward velocity) in between, that is when sweeping past its general direction of travel. Outside of this functional range, regular oscillations disappear for extreme parameter sets. For instance, if one neuron overpowers the other due to too strong reciprocal inhibitory strength (C, overpowering); or inversely, due to the absence of reciprocal inhibition (strength <0.01) or a too strong internal adaptation preventing the neurons to modulate their baseline activity, and thus resulting in noise (A).

## Discussion

### An internally generated movement at the core of ant visual navigation

We recorded the detailed locomotor movements of two distantly related ant species – *M. croslandi* and *I. purpureus –* using a trackball-treadmill device directly in their natural environment. Both ant species displayed a similar rhythmical pattern of movement combining continuous lateral oscillations with a synchronized variation of forward speed: ants slow down – sometimes until a complete halt (see video Fig. 4D) – at the end of each left and right sweep, that is, when their body is facing away from their overall direction of travel. Conversely, they display a burst of forward speed in between, that is, when their body is aligned with their overall direction of travel, even though during these moments angular speed is rather high (Figs. 3, 5A, S3). This generates oscillatory trajectories that optimizes the distance covered when facing the general direction of travel *–* and thus the effective distance covered *–* while simultaneously scanning left and right directions.

This particular movement signature is conserved whether ants are on their familiar route, in an unfamiliar panorama (Fig. 3) or when displaying their natural learning walks around the nest (Fig. 5A). Also, this movement signature is still expressed when the ant body is artificially fixed on our trackball in a given orientation (Fig. 6; Fig. S3), which shows that it results from an internal process rather than a servomechanism based on external cues. Previous observations assumed that ants accelerate when facing their goal direction as the response to the recognition of familiar views when aligned in this particular direction (Baddeley et al., 2012; Kamhi et al., 2020; Wystrach et al., 2013). Here we show that this acceleration is a product of an endogenous process (that is also expressed in unfamiliar terrain), and thus stresses the often disregarded importance of internally generated movement within the ‘sensorimotor’ loop (Brembs, 2021; Gomez-Marin et al., 2011; Schroeder et al., 2010; Wolf et al., 2017; Yuste et al., 2005).

### Modulation of the exploration/exploitation balance

The described internally generated movement dynamic provides an embedded solution for the compromise between exploration and exploitation. External visual cues can then modulate the exploration/exploitation balance to the task at hand by simply adjusting the amplitude of this endogenous dynamic. Higher amplitude oscillations optimize exploration in an unfamiliar environment, while lower amplitudes favor ‘exploitation’ (straighter paths) on a familiar route. Interestingly, this movement signature is equally useful, and used, during the acquisition of learnt visual information, when naïve ants display their learning walk around the nest (Jayatilaka et al., 2018) (Fig. 5). Optimising the acquisition of information requires to bias the balance toward ‘exploration’ in the same manner as searching for familiar cues in an unfamiliar environment does; thus it is not surprising to see here also high amplitude oscillations (Fig. 4, 3). The apparent different needs for the acquisition of information on the one hand and the use of that information on the other hand, are thus solved with the same solution. The weaker modulation observed in Iridiomyrmex may be explained by their strong use of chemical trails (Carde et al., 2016), whose absence on the trackball may favor exploration in both visually familiar and unfamiliar conditions.

Finally, we see in both species that the internally generated oscillations can be strongly – if not entirely – inhibited when not needed, such as when moving in the dark (Fig. 2). Indeed, oscillations are not always useful, and it may be advantageous to repress them in situations like when inside the nest, or when trying to escape an aversive situation. It is thus expected that the animal’s internal state as well as the presence/absence of relevant contextual information – such as a visual panorama by opposition to no panorama at all – modulates the expression of the internally produced oscillations in a similar vein as the male fly’s internal state continuously tunes up or down the gain of circuits promoting visual pursuit (Hindmarsh Sten et al., 2021). However, what specific type of visual information promotes the expression of oscillations remains to be seen.

### Neural Implementation

In the insect brain, regular lateral oscillations seem to result from reciprocal contra-lateral inhibitions between left and right hemispheric premotor areas, so-called Lateral Accessory Lobes (LAL). These circuit results in the internal production of asymmetrical and rhythmical excitation between left and right motor commands, so called flip-flop neurons (Berni, 2015; Berni et al., 2012; Iwano et al., 2010; Kanzaki, 2005; Namiki and Kanzaki, 2016b; Steinbeck et al., 2020). Lateral oscillations are thus generated endogenously by structure analogous to central generator patterns (CPG). CPGs sustain a wide range of rhythmic functions such as chewing, breathing or various locomotor movements and can be typically modulated by sensory input (Marder and Calabrese, 1996; McAuley et al., 1999; Schroeder et al., 2010; Wolf et al., 2017; Yuste et al., 2005). Here however, rather than limb movements, the CPG-like in the insects’ LAL controls the displacement of the whole animal across space and thus, directly contributes to the navigational task.

The systematic positive covariation between angular and forward velocity observed here (Figs. 3, 5, S3) are unlikely to result from physical body constraints (where one could expect the opposite, that is, a negative correlation between angular and forward velocities) and thus suggests that the internal dynamics of the LAL control both turning and forward velocity simultaneously. Remarkably, the observed relationship between forward and angular velocity readily emerges from our toy model when simply assuming that forward speed results from the overall excitation across both sides of the LAL; while turning velocities results from the difference between left and right excitation (Fig. 8). Bursts of forward speed appear when one side largely dominates over the other, that is when the ant is at the maximum speed of its angular sweep and thus roughly aligned with its overall direction of travel. Inversely, a break of forward speed occurs when the dominancy is reverted between LAL neurons, that is at the moment where ant reverts their turning direction (Fig. 8), as we can see in our ant data (Fig. 3, 4, 5, S3). Certainly, we do not exclude that more complex circuitry could produce other regimes of covariations but our simple model shows that this particular relationship between forward and angular speed can readily emerge and is robust to modulation of the system. In contrast, the oscillations’ amplitude and frequencies are somewhat sensitive to parameter change, at least across a factor of 2 (Fig. 8—figure supplements 1). Therefore, we can easily envision how various inputs into the LAL (such as pictured in Fig. 9) could modulate the amplitude and frequency of the emerging oscillations as we observed in our various conditions, without altering the fundamental relationship between forward and angular velocities. Understanding how such inputs actually operate in insects opens the door to a complexity that would be too vast to tackle here, but given the rapid development of neurobiological tools and knowledge about the circuitry of the lateral accessory lobes, this may constitute a realistic endeavour in the near future.

**Figure 9.**
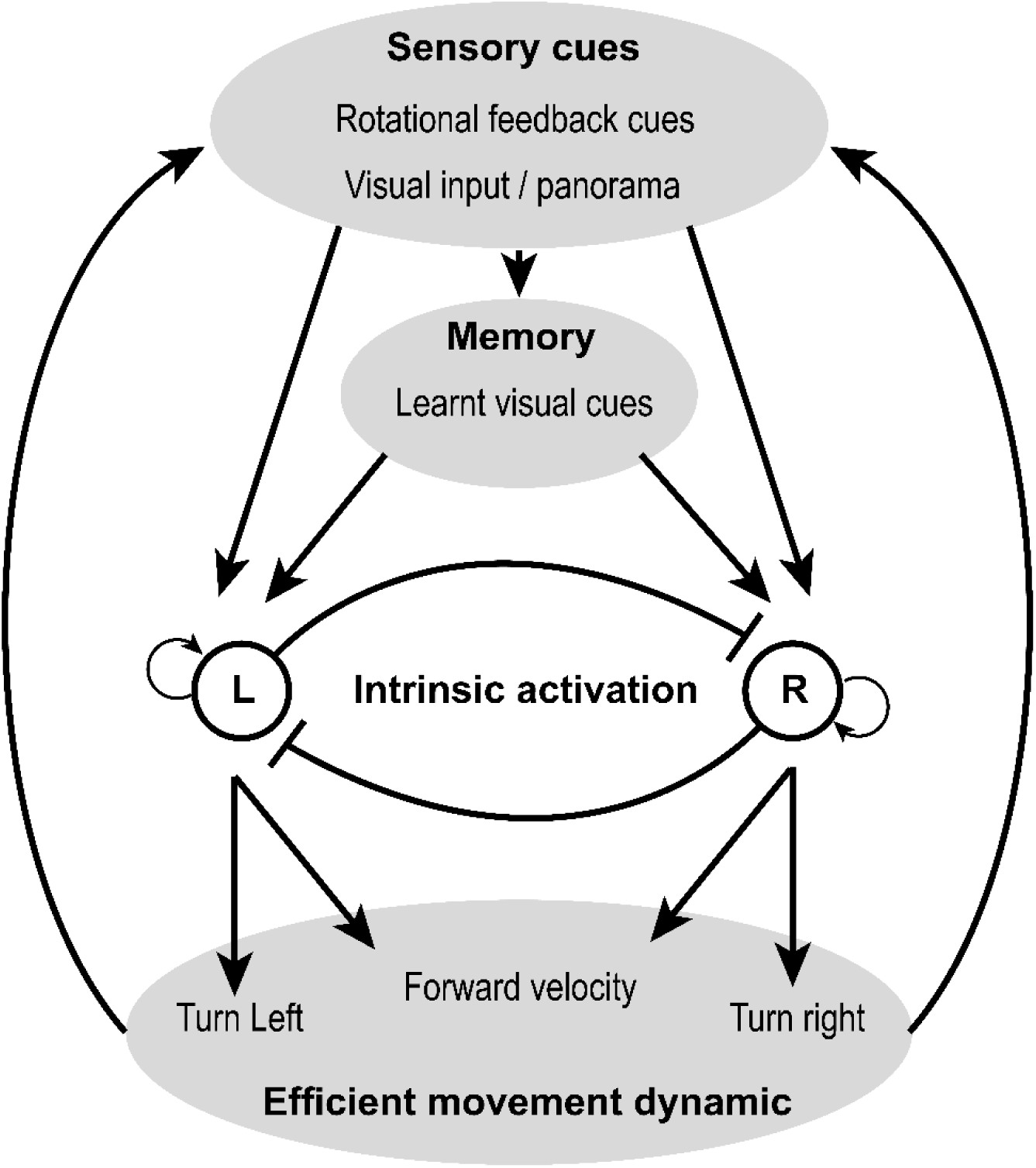
An intrinsic oscillator at the core of visual navigation. We propose a simple scheme to encompass our results. In this view, sensory input (in this case various innate and learnt visual information) act on behavior only indirectly, through the systematic modulation of an intrinsic oscillator. The latter ensures that an efficient movement dynamic is preserved across navigational contexts, which would not necessarily be the case if various and potentially conflicting sensory information were directly modulating movement. This scheme also highlights the idea that action is not merely the product of perception. At the core of behavior lies an intrinsic, self-generated dynamic, which are modulated, rather than controlled, by sensory perception.

### Evolutionary consideration

Several clues point to the idea that displaying such oscillations is an ancestral way of moving, that predates at least the holometabolous insects’ common ancestor (around 300 Myr ago). Indeed, comparable oscillations around 0.3-1Hz are observed in the two ant species studied here but also in moths (Kanzaki et al., 1992; Kanzaki and Mishima, 1996; Kuenen and Baker, 1983; Olberg, 1983) *Drosophila* larvae (Wystrach et al., 2016), Colorado beetles (Lonnendonker, 1991) and other insects (Namiki and Kanzaki, 2016a). Furthermore, the LAL that likely produces these oscillations is an ancestral brain structure, largely conserved across insects (Namiki and Kanzaki, 2016b, 2016a; Steinbeck et al., 2020).

Some have suggested that neural systems evolved at first to coordinate movements endogenously (Keijzer et al., 2013; Yuste et al., 2005) whereas the modulation of these movements by external sensory information originated only as a second step (Keijzer et al., 2013). This view fits the observation that various multi-modal sensory cue converge to the same conserved region (the LAL) to produce a remarkable variety of navigational behaviors (Namiki et al., 2014; Namiki and Kanzaki, 2016b; Steinbeck et al., 2020). In crawling *Drosophila* larvae oscillations are modulated by variations in odor concentration, resulting in attraction toward (or repulsion from) the odor source (Wystrach et al., 2016). In flying male moths, it is the detection of a female’s pheromone that modulates oscillations and enables the insect to track the odor plume. In walking ants also, zigzagging paths seem to be modulated by olfactory cues when following chemical trails (Hangartner, 1967). In wasps, the presence of a proximal visual object appears to trigger zigzag flights, which may provide useful visual depth information to the insect (Voss and Zeil, 1998). And finally, in visually navigating ants, oscillations are modulated by the recognition of familiar views to support route-following, visual exploration and the acquisition of visual memories in naïve individuals.

The modulation of oscillations by rotational visual cues, as we showed here in ants (Figs. 6, 7) might be ancestral to insects as we equally observe it in moths (Namiki et al., 2014; Pansopha et al., 2014) (Namiki et al., 2014; Pansopha et al., 2014). This likely evolved through connections between horizontal optic-flow detectors in the visual lobes (Busch et al., 2018) and the LAL (Fig. 9), providing a useful feedback-control to calibrate the amplitude of turns, and more generally participate in the widespread optomotor response. The modulation of oscillations by the presence/absence of visual cues, and more particularly familiar/unfamiliar visual cues however, must have appeared later with the evolution of visual route-following and homing in hymenopteran, through direct or indirect connections between the Mushroom bodies – the seat of navigational visual memories (Buehlmann et al., 2020; Kamhi et al., 2020; Webb and Wystrach, 2016) – and the LAL (Fig. 9).

Finally, the forward speed variation within the oscillation cycles presented here – which can emerge readily from the LAL connectivity (Fig. 8) – has not been reported in other species and thus could be a derived adaptation in ants. This may be an adaptation for walking. Even though ants can decouple their body orientation from their direction of travel if needed (Schwarz et al., 2017), walking sideways may be inefficient and preferably avoided due to physical constraints on leg movements. Thus, slowing down when looking on the side may be a compromise between the preference for walking forward (rather than sideways) and avoiding drifting away from the desired travel direction. Alternatively, this dynamic may be an adaptation for visual cue recognition in navigation, by opposition to the use of olfactory cues. When tracking an odor plume or trail, movements off track to the left and right (and crossing in between), provide useful information regarding the direction to follow. However, such sideway displacement off the beeline become useless for visual recognition (gaze *orientation* is what matters (Zeil 2012; Wystrach 2021)) and thus costly as they increase the overall distance walked. In addition, pausing when looking on the side must improve the efficiency of visual recognition due to gaze stabilization, which accommodates remarkably well the idea that views are learnt and recognized when oriented left and right, rather than toward and away from the goal (Stürzl et al., 2016; Wystrach et al., 2020).

These two hypotheses (i.e., adaptation to walking vs. visual recognition) could be tested straightforwardly by investigating the existence of such burst of forward speed during route-following in (1) walking ant species that do not use vision or in (2) flying insects that do use vision and are known to oscillate (e.g., bees or wasps) (Egelhaaf et al., 2012; Stürzl et al., 2016; Voss and Zeil, 1998). Overall, it is remarkable how a basal endogenous circuit producing a rhythm can be exapted to receive such a diversity of modalities and produce such a diversity of behaviours (Wystrach, 2021).

## Conclusion

Tracking a pheromone plume using olfaction, walking along a route using learnt visual cues or performing a learning walk as a naïve ant may appear as quite different tasks. However, our results support the idea that the solution to these challenges is based on a shared mechanisms: namely, the intrinsic production of left/right oscillations in a conserved pre-motor area (Kanzaki et al., 1992; Kodzhabashev and Mangan, 2015; Le Möel and Wystrach, 2020; Murray et al., 2020; Pansopha et al., 2014; Steinbeck et al., 2020). This internally generated movement dynamic provides a remarkably optimized solution to the compromise between exploration and exploitation. Various sensory cues, through both innate or learnt pathways, can alter the exploration/exploitation balance to fit the task at hand by simply adjusting the amplitude of this endogenous dynamic. Evolution appears to have favored a general-purpose steering mechanism and it is notable how such a simple and repetitive oscillatory movement, generated intrinsically, accommodates such a diversity of input and be useful for such a diversity of tasks across species, modalities and ways of moving (i.e., crawling, flying or walking).

## Methodology

### Study Animal & Experimental site

All experiments took place within an open grassy woodland at the National Australian University, Canberra from Feb. to Mar. 2019. We had the opportunity to work with two Australian endemic ant species: *Myrmecia croslandi* and *Iridomyrmex purpureus. Myrmecia croslandi* workers are known to forage solitarily, with each individual either hunting on the ground in the vicinity of the nest or navigating routinely back and forth toward the same favorite foraging tree throughout her life span (Jayatilaka, 2014). These ants rely mainly on learnt terrestrial visual cues to navigate but are also able to resort to PI when the visual environment does not provide guidance (Jayatilaka et al., 2014, 2018; Narendra et al., 2013; Zeil et al 2014). Eight nests of *M. croslandi*, that foraged on two distinct trees between 6.0 and 30.0 m away from the colony, were used in this study. *Iridomyrmex purpureus* ants form large colonies and forage along pheromone trails that lead to food patches. Despite employing pheromone trails for recruitment, this species also uses both learnt visual information and path integration for navigation (Card et al., 2016). We also performed navigational experiments with two *I. purpureus* nests to ensure that this species relies indeed on both learnt visual terrestrial cues and path integration, which they did surprisingly well (see Fig. S1). For the main experiment, a feeder was placed 7.0 m away from the nest and foragers were free to familiarize themselves with the route for 72 h before being tested.

### Trackball system and data extraction

During tests, ants were mounted on a trackball device (Dahmen et al., 2017). This device consists of a polystyrene ball held in levitation in an aluminum cup by an air flow. The trackball has two sensors placed at 90° to the azimuth of the sphere, which record the movements of this sphere and translate them into X and Y data retracing the path of the ant. The X and Y acquisition of the trackball rotations happened at a 30 Hz frequency (i.e., 30 data points per second), enabling us to reconstruct the ant’s movements with high precision. Additionally, a camera (640×480 pixels) recording from above provided details of the ant’s body orientation, also at 30 Hz. We also analyzed *M. croslandi* learning walks recorded directly on the natural ground. Head directions were obtained via video recordings at 25 Hz (provided by Jochen Zeil).

In this study, we used two different trackball configurations to record the ants’ motor responses: further referred to as ‘free ant’ and ‘fixed ant’ experiments, respectively. Free ant experiment: two small wheels prevented the polystyrene ball from rotating in the horizontal plane, however, all other degrees of freedom of the ball rotation were accessible (Fig. S1 A; ‘closed-loop’ in Dahmen et al., 2017). Ants were attached on top of the ball by putting magnetic paint on their thorax and a micro-magnet fixed at the bottom end of a single dental thread that was in turn attached to a 0. 5 mm pin. Crucially, the pin was placed within a glass capillary. This procedure enabled the ants to execute physical rotations on the ball (the ball is not rotating horizontally) but prevented any translational movement. Ants could thus execute body rotations and control the direction in which they faced but any attempt to go forward or backward resulted in ball rotations.

Fixed ant experiment: the two small wheels were no longer in place. Hence, the ball could now turn in any direction, including the horizontal plane (Fig. S1B; ‘open-loop’ in Dahmen et al., 2017). Ants were tethered directly to a needle with a micro magnet (glued at the end of the needle) and magnetic paint on their thorax. The top end of the needle was glued to a small piece of paper sheet (0.5×2.5 cm). Consequently, the fixed ant could no longer rotate, and the experimenter could choose in which direction she was facing. Any attempt to move, including turning, by the ants resulted in ball rotations. This trackball configuration was only conducted with the more robust *M. croslandi*.

### Free ant experiments: protocol

At the start of each test, the ant was mounted on the trackball device but surrounded by an opaque cylinder (30×30 cm) that prevented the ant from seeing any cues from the visual scenery around her. Once the ant was in place on the device, the whole apparatus was moved to the desired test location in the field. Afterward, the surrounding cylinder was removed, revealing the visual scenery to the ant and data recording began. To ensure a high level of homing motivation, only individuals who had previously received a 40% sucrose solution (for *M. croslandi*) or a food item (for *I. purpureus*) were tested. The recording period was 3.5 min for the robust *M. croslandi* and 1.5 min for the flimsier *I. purpureus*.

To test the impact of terrestrial visual cues on the oscillation behavior, ants were tested under three distinct conditions.

Familiar (F): ants were tested along their habitual route, which therefore presents a familiar visual view.

Unfamiliar (U): ants were tested at least 50 m away from the habitual route, which therefore presents an unfamiliar panorama.

Dark (D): ants were tested in total darkness, within an opaque cylinder (30×30 cm) covered with a red Plexiglas plate that transmitted only the low red wavelengths, which ants cannot perceive (Aksoy and Camlitepe, 2018; Ogawa et al., 2015).

To test the impact of PI on oscillation behavior, ants were tested as either full- or zero vector ants. Full vector (FV): ants were caught at the start of their inbound trip to the nest (i.e., at the foraging tree for *M. croslandi* and at the feeder for *I. purpureus*). FV ants have an informative homing PI vector, which points in the food-to-nest compass direction. Consequently, FV ants can rely on both PI vector and the learnt visual scenery while being tested. Zero vector (ZV): homing ants were captured just before entering the nest, that is, at the end of their inbound trip. Hence, their PI homing vector is reduced to zero and thus no longer directionally informative. Consequently, ZV ants can only rely on the learnt visual scenery while being tested. For each of the three visual conditions FV and ZV ants were tested, resulting in a total of six conditions.

For *I. purpureus*, at least 16 ants were tested in each of the six conditions (F: FV&ZV =16; U: FV&ZV =17; D: FV=16 & ZV=17). Since *I. purpureus* forms very populous colonies with an abundance of foragers, all individuals were tested only once, in one of the six conditions. The data obtained are therefore statistically independent. On the contrary, *M. croslandi* forms sparsely populated colonies and individuals usually make only one foraging trip per day (Jayatilaka 2014, a, b). Thus, it is time-consuming and challenging to capture, mark and follow individuals throughout foraging trips. Individuals were therefore captured (either at the foraging tree (FV) or before reaching their nest (ZV)) and tested successively in a pseudo-random order in all three visual panoramas: Familiar (F), Unfamiliar (U) and in the Dark (D). Both the sequence in which the visual conditions were experienced and the individuality were included in the statistical models as it is likely that the state of the PI vector will be modified across successive tests. Overall, 32 *M. croslandi* ants were tested with some individuals tested as ZV or FV ants on two different days. At least 16 *M. croslandi* ants were tested in each of the six conditions (F: FV&ZV =17; U: FV&ZV =17; D: FV=17 & ZV =16). Overall, 101 recordings were obtained.

In the ‘free ant’ experiments, the ant’s body axis can turn without the ball movement. We used the recorded videos to manually track the ants’ body orientation through time using the free software Kinovea. We removed the first 3s of recording of each ant as the removal of the opaque ring may have disrupted the behavior. Overall, the analyzed recording length was 100s (except for two ants: 84 and 99s) for *M. croslandi* individuals and 50s (except for two ants: 49s) for *I. purpureus* ants. For details of the Fourier analysis (see below); nine paths of *I. purpureus* were discarded (F FV=1, U FV = 1, U ZV =2, D FV =2, D ZV = 3) as the ants displayed too many pauses.

### Fixed ant experiments: protocol

To test if oscillations are due to an intrinsic oscillator or caused by the fact that ants try to keep a bearing toward an external stimulus, we conducted an additional experiment on *M. croslandi* ants. We recorded foragers without an informative integration vector (ZV) in the same unfamiliar environment (U) as before. But this time, ants were tested in eight different orientations, covering the 360° azimuth by bins of 45°; and with one direction corresponding to the food-to-nest compass direction. Each ant was tested in all eight orientations in a pseudo-random sequence. To change the ant’s orientation, the experimenter would first place the opaque cylinder (30×30 cm) around the trackball to prevent individuals from perceiving any visual panorama, then rotate the whole set-up (trackball and mounted ant) and finally remove the opaque cylinder to re-start data collection. Ants were recorded for at least 15s up to 20s in each orientation. Eleven ants were tested in all eight orientations. Since the ants were fixed, trackball rotations along the horizontal axis provided a direct measure of the angular velocity of the attempted turn generated by the ant. Angular velocity time series were then extracted from the trackball data. At the end, the available length of the recordings for the analysis had 512 frames (∼17s) except for 5 individuals (294,394,456,474,494 frames). It should be noted here that prior to the Fourier analysis (see below) we added a series of 0s (zero padding) at the end of the time series until it had a length of 3000 data points to match the same recording length obtained in the free ant experiment). This permits to increases the precision in the frequency range obtained from the Fourier analysis.

### Fourier analysis

To reveal the occurrence of regular lateral oscillations, we choose to focus on the angular velocity value (Fig. S1C), which constitutes a direct reading from the left/right motor control. This time series was processed through three successive steps to obtain its ‘spectral density’, according to the Wiener-Khinchin theorem. First, the signal was parsimoniously smoothed with a moving median running of a length of five frames (0.17s) to reduce the influence of the recording noise (Fig. S1D, dashed blue line). Then, the recorded time series was passed through an autocorrelation function (Fig. S1D). Finally, a Fourier transform was performed on these autocorrelation coefficients, providing the power spectral density (Fig. S1E). With this approach, the magnitudes obtained are independent from angular amplitudes of the oscillations and thus can be directly compared across individuals. A high magnitude indicates a strong oscillation for a given frequency. For each individual, the dominant frequency (i.e., presenting the highest peak magnitude) and its magnitude were extracted (Fig S1E, dashed blue lines). To check whether these magnitudes indicate a significant regular oscillation, we compared them to the average spectral density magnitude resulting from 200 Gaussian white noise signals of the same length (within each species and experiment). Gaussian white noise signals were obtained by drawing a sequence of random values drawn from a normal law. These simulated signals were then processed through the exact same operations as the ants’ angular velocity recording: namely smoothing, autocorrelation, Fourier transformation and extraction of the highest peak magnitude. The mean of the 200 highest magnitudes obtained was then compared to the real ants’ equivalent magnitudes with a Wilcoxon one-tail test. As each experimental group has been compared with this mean magnitude of the simulated angular velocity signal, the p-value is subsequently adjusted using the Bonferroni correction.

### Average oscillatory cycle

To extract the average dynamics of an oscillation cycle, the mean cycles at the individual and population levels were reconstructed as follows. First, we smoothed the angular velocity time series of each individual by running twice a median with a window length of 31 frames for *M. croslandi* (for both free and fixed experiments), 25 frames for data during learning walks on the natural ground and seven frames for *I. purpureus*. This window length is much smaller than one oscillatory cycle and thus smoothens the data without altering the cycle general dynamics (Fig. *3*—figure supplements 1A). Second, we indexed moments when the time series crosses 0 (from – to +) as t_0_; and extracted a window of ±90 frames around the t_0_ indices (±60 frames for *I. purpureus*; Fig. *3*—figure supplements 1A). We then reconstructed a mean cycle for each individual by averaging the individual’s extracted windows, aligned at t_0_ (Fig. *3*—figure supplements 1B). The individuals’ average forward speed dynamics during one cycle was obtained in the same way by using the same indices t_0_ obtained from the angular velocity data (Fig. *3*—figure supplements 1C). For each ant, the mean angular velocity peak and amplitude of forward velocity cycles were extracted for analysis. Finally, we reconstructed the average cycle at the level of the population by averaging the mean cycle of all individuals. Note that to do so, each individual’s mean cycle was first normalized to show similar amplitudes (mean_cycle_normalised = mean_cycle/mean|mean_cycle_values|). The goal was here to observe the cycle dynamics through time and not to estimate the inter-individual variation in amplitude, which was analyzed previously using the individual’s mean cycle amplitude. Note that the criterion used to align the time series (the change from a left to a right turn) necessarily creates an artefact in the averaged angular velocity obtained. Namely, an average change from left turn to right turn at t_0_. However, several factors indicate the relevance of such pooling at the population level: (1) the period of the average cycle corresponds to the mean frequency obtained from the Fourier transform. (2) we can observe a significant reversal of the angular velocity before and after the mean oscillation cycle. (3) the associated forward velocities co-vary in a significant way, indicating the existence of conserved dynamics.

### Analysis of the ants’ direction of movement

We reconstructed the ant paths that derived from the trackball recording to determine the mean direction of movement (mu) as well as the mean circular vector length (r, a measure of dispersion) of each individual. The mean directions (mu) were analyzed using a Rayleigh test (from R package: Circular) that also includes a theoretical direction (analogous to the V-test). To test whether the angular data are distributed uniformly as a null hypothesis or if they are oriented toward the theoretical direction of the nest as indicated by the state of the PI or the familiar panorama, the average vectors lengths (r) were analyzed via a Wilcoxon-Mann-Whitney test with a Bonferroni correction for multiple testing.

### Model

For the free ant experiment, two types of models were tested: the first considering the interaction between visual panorama and vector modalities, the second with simple additive effect. It should be noted here that in case of deviation of the residuals of these models from normality and or homoscedasticity, the response variables have been transformed. Only the model presenting the lowest Akaike information criterion was retained and subsequently analyzed via an analysis of variance (Anova), followed by a post-hoc analysis of Tukey’s rank comparison. Finally, for *M. croslandi* whose individuals were tested several times, the models are mixed models that consider these two random variables. For some models, we removed the sequence effect because of a singularity problem of this factor (i.e., the information given by this variable is already contained in other variables). For fixed ant experiments (which were tested only as ZV ants in an unfamiliar environment), the model analyzed the effect of the condition of the subsequent rotations. All data were processed and analyzed using free software R version 3.6.2.

## Acknowledgments

We want to thank the Australian National University and particularly Jochen Zeil, Zoltán Kócsi and Trevor Murray for their advice and technical support. We also thank Hansjürgen Dahmen for providing us with the trackball system and Ajay Narendra for his incredible knowledge about the local Australian ants. We are grateful to all these people for their helpful advice. We also thank Jochen Zeil, Paul Graham and Michael Mangan for their helpful feedback on the manuscript. Finally, we thank ants tested in these experiments for their participation.

## Additional information

### Author contributions

LC Conception and Design of experiment, Data collection, Analysis and interpretation of data, Drafting and revising the article. SS Conception and Design of experiment, Data collection, Drafting and revising the article. AW Conception and Design of experiment, Data collection, Analysis and interpretation of data, Drafting and revising the article, Supervision of project.

## Supplemental information 1

To this date, the navigational skills of *Iridomyrmex purpureus* have been investigated in only one study showing that these individuals are able to use terrestrial as well as celestial cues for orientation (Card et al 2016). Before using this species in our study, we performed similar experiments and found that this ant relies indeed on both sources of information to navigate. First, to test whether *I. purpureus* was able to use visual familiarity foragers were capured at the end of their inbound trip where the vector of integration was reduced to zero (ZV) and re-released (tested) along their route. Second, to test if *I. purpureus* was able to use celestial information foragers were captured at the feeder at the start of their inbound trip (FV) and re-released (tested) in an unfamiliar location ∼50 m away (see Methodology).

For tests, ants with a food item were released at the center of a goniometer divided in 18 sectors (20° each) either in a familiar or unfamiliar surroudings. The taken directions of the tested ants were recorded at two points: 15 cm and 27 cm. We used the “corrected direction” for our analyses, which is the direction taken between these two points. Note, that after several tested individuals, the goniometers was rotate (180°) to prevent the use of chemichal trails.

**Figure S1:**
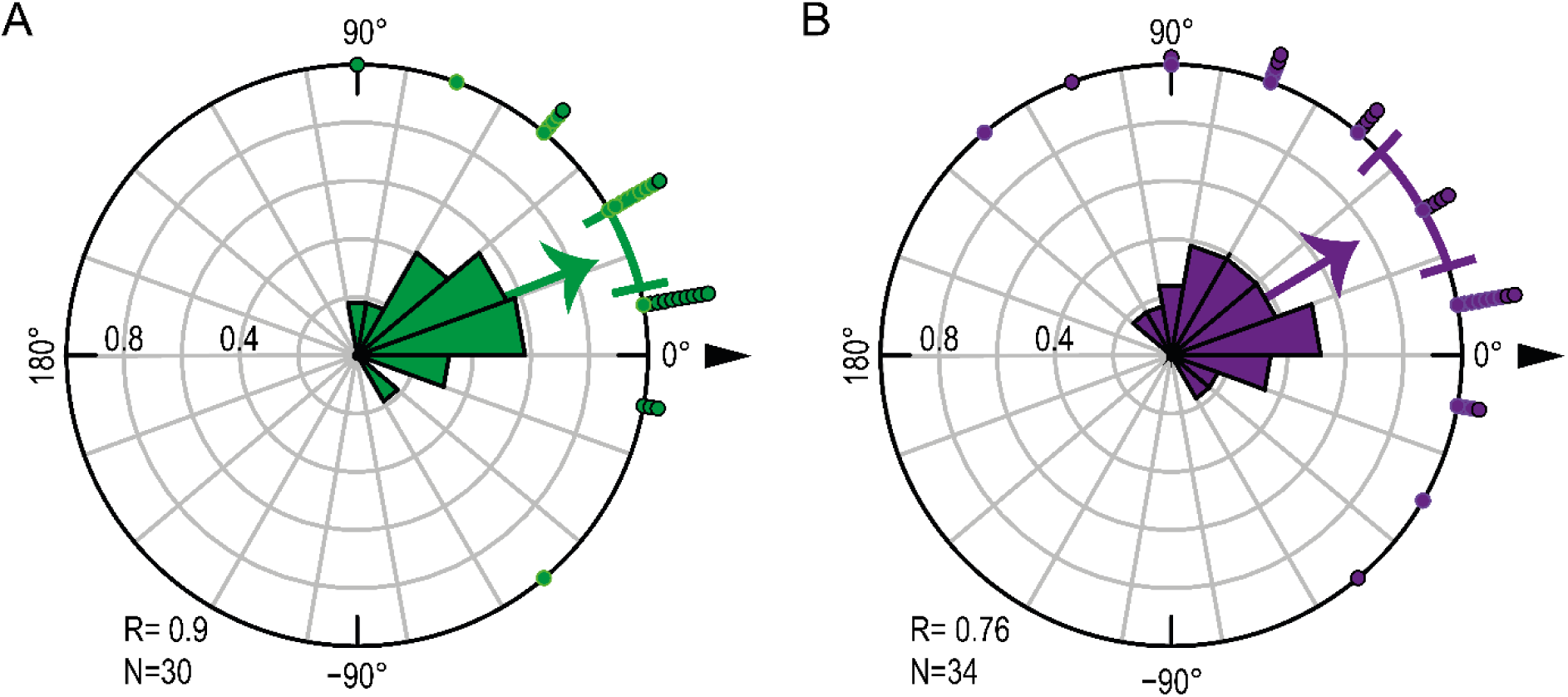
The meat ant *Iridomyrmex purpureus* uses both visual and celestial cues to navigate. (A) Circular histogram depicting the directional decisions of tested zero-vector ants (without informative integration vector). (B) Circular histogram depicting the directional decisions of tested full-vector ants (with informative integration vector). For A and B, nest direction is indicated by the 0° direction (black triangle). The arrow corresponds to the mean vector of the distribution. The 95% confidence interval of the mean is displayed as a colored arc. Each dot corresponds to one corrected direction obtained from one individual. Dots with a black rim are data recorded before rotation of the goniometers and the dots without black rim after rotation.

First, the results show that ZV ants of *I. purpureus* use the familiar visual surrounding to orient toward their nest (Fig. S1A; Rayleigh test with nest as theoretical direction 0°; p<0.001). Secondly, in unfamiliar panorama results clearly show that FV ants followed the direction indicated by celestial cues (0° on the Fig. S1B; Rayleigh test p<0.001). Therefore, this species does not only rely on chemical trails but also on the visual panorama and the path integration during foraging.

## Supplemental information 2

**Figure S2:**
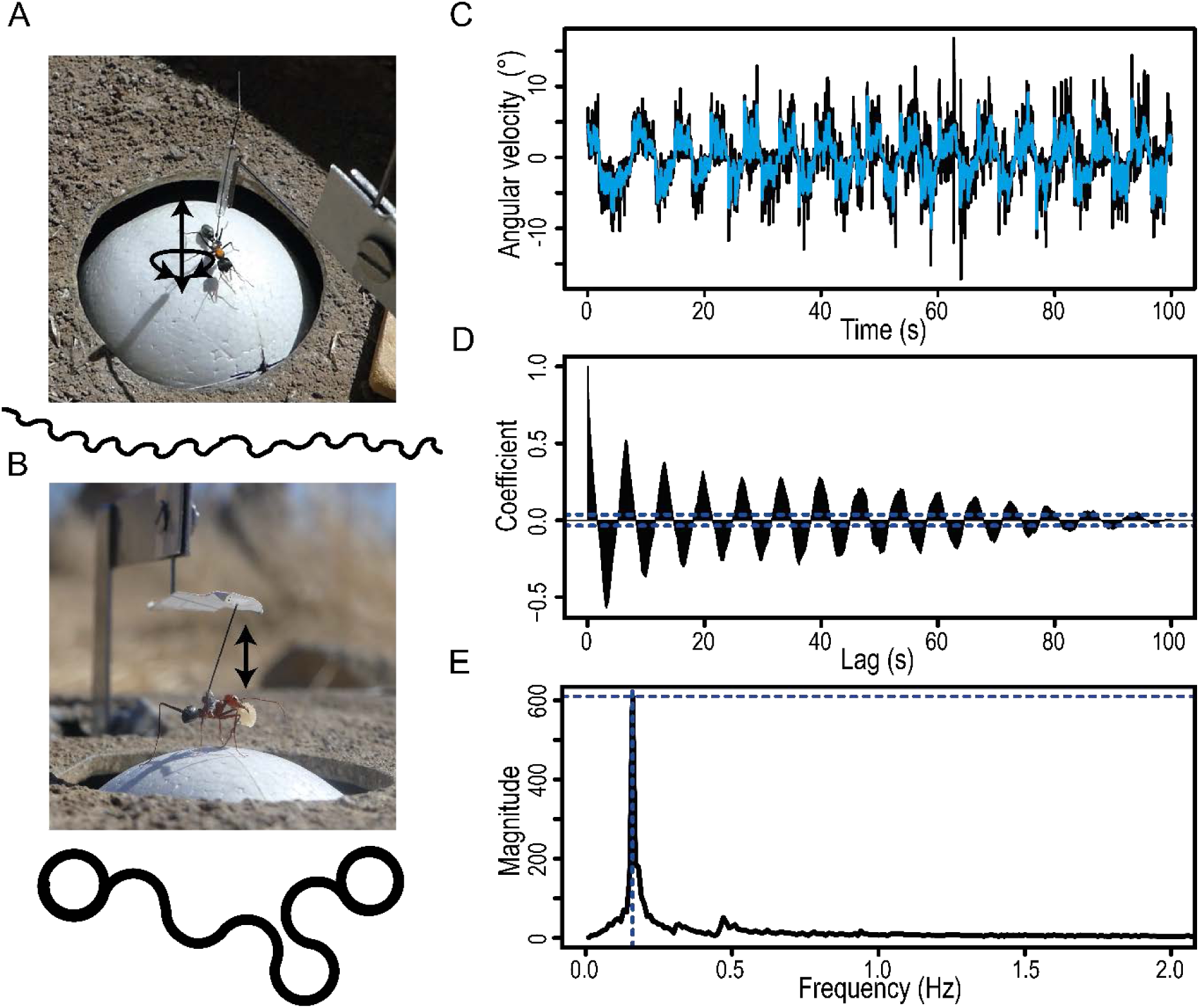
Trackball set-ups, recording and processing of the ants’ trajectory. (A) Trackball set-up for ‘free ant experiment’ (top view). Two wheels prevent the sphere from rotating along the horizontal plane. Ants are free to rotate their body along the yaw axis to control in which direction they perceive the world. (B) Trackball set-up for ‘fixed ant experiment’ (side view). The two wheels have been removed, so the ball can turn in every direction. Ants are fixed in a way that prevent body rotations along the yaw axis, so they are forced to keep their bearing in the imposed direction. For A and B an example path has been plotted below the picture. (C) Angular velocity signal over time (recorded at 30 readings/sec) in an individual of the species *M. croslandi* (black) with smoothed signal superimposed (blue). (D) Autocorrelation carried out on the entire smoothed angular velocity signal. (E) Fourier transformation of the autocorrelation coefficients signal (shown in D) provides the ‘spectral density’. This approach has the advantage to provide magnitudes that are directly comparable between individuals. For each individual, the frequency peak with the highest magnitude was extracted, indicating a strong oscillation of the angular velocity signal at that frequency (dashed blue lines).

## Supplemental information3

**Figure S3:**
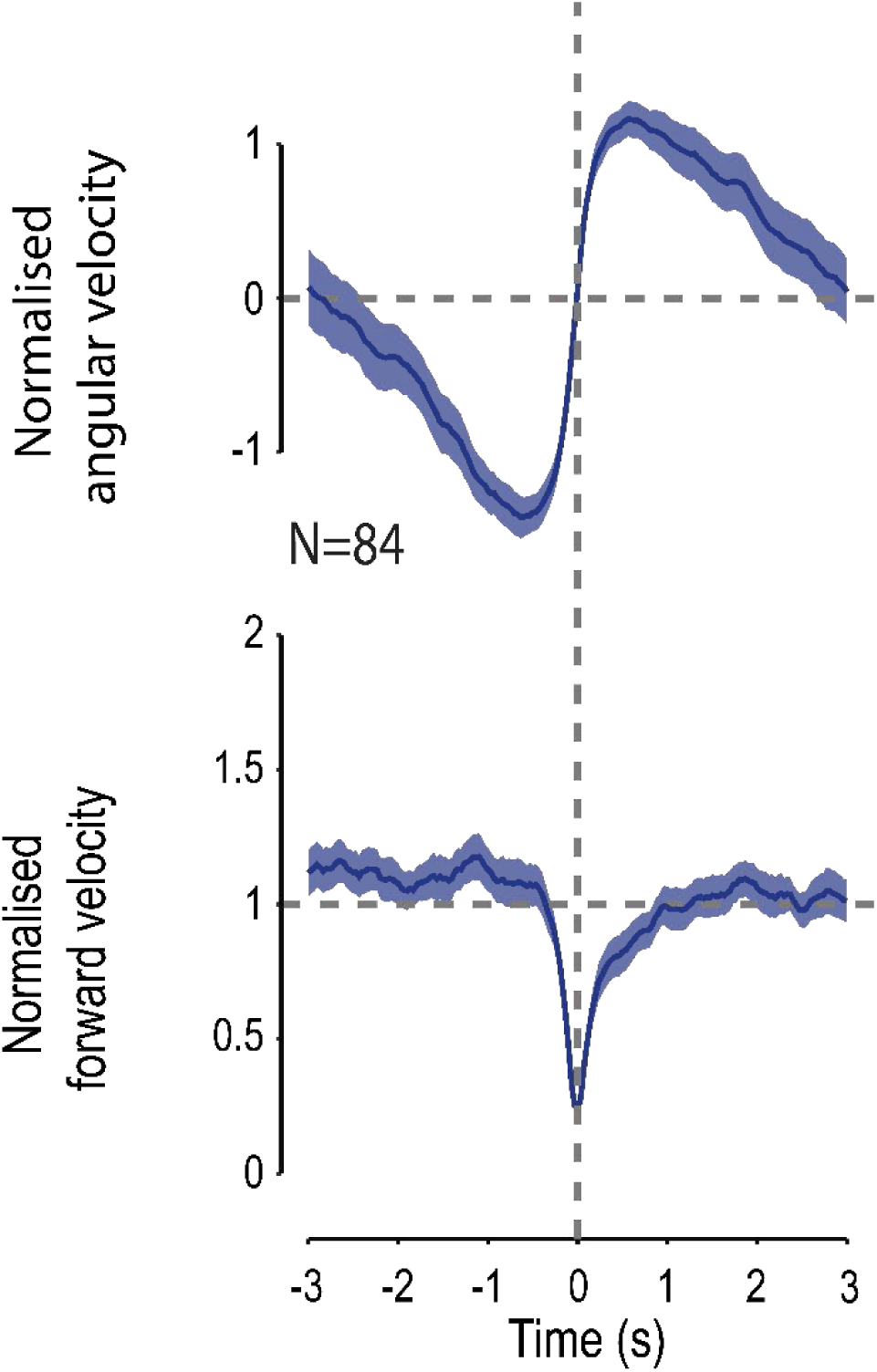
Absence of the rotational visual feedback does not modify the dynamical signature of oscillation behavior. Angular velocities (top-row) and forward velocity (bottom row) co-varies in a way that seem conserved across experimental conditions (Fig. 3 & Fig. 5). Population cycles have been reconstructed by merging all rotational condition as there is no differences of amplitude of mean peak of angular velocity and mean forward amplitude cycles between rotational conditions (Anova: Df= F_7,84_= 1.0103, p≥0.432). Each individual signal has been normalized before pooling. Colored areas around mean curves represent the 95% confidence interval, based on the inter-individual variation.

